# QAlign: Aligning nanopore reads accurately using current-level modeling

**DOI:** 10.1101/862813

**Authors:** Dhaivat Joshi, Shunfu Mao, Sreeram Kannan, Suhas Diggavi

## Abstract

**Motivation:** Efficient and accurate alignment of DNA / RNA sequence reads to each other or to a reference genome / transcriptome is an important problem in genomic analysis. Nanopore sequencing has emerged as a major sequencing technology and many long-read aligners have been designed for aligning nanopore reads. However, the high error rate makes accurate and efficient alignment difficult. Utilizing the noise and error characteristics inherent in the sequencing process properly can play a vital role in constructing a robust aligner. In this paper, we design QAlign, a pre-processor that can be used with any long-read aligner for aligning long reads to a genome / transcriptome or to other long reads. The key idea in QAlign is to convert the nucleotide reads into discretized current levels that capture the error modes of the nanopore sequencer before running it through a sequence aligner.

**Results:** We show that QAlign is able to improve alignment rates from around 80% up to 90% with nanopore reads when aligning to the genome. We also show that QAlign improves the average overlap quality by 9.2%, 2.5% and 10.8% in three real datasets for read-to-read alignment. Read-to-transcriptome alignment rates are improved from 51.6% to 75.4% and 82.6% to 90% in two real datasets.

**Availability:** https://github.com/joshidhaivat/QAlign.git

## 1 Introduction

In genomic data analysis, aligning DNA / RNA-seq reads to a genome / transcriptome is a key primitive, that precedes many downstream tasks, including genome / transcriptome assembly [1, 2] and variant calling [3, 4, 5]. Getting accurate read alignment is difficult especially in repetitive regions of the genome, due to the short length of the reads obtained via high throughput sequencing. Emerging sequencing technologies, particularly, nanopore sequencing [6, 7] offers a potential solution to this problem by providing long reads (with average read length 10-kb and the longest read sequenced so far *>* 2Mb) that can span these repetitive regions. However, these long reads are riddled with a high error rate, thus, making alignment of low accuracy [8] and the downstream task difficult. For example, while nanopore sequencing has enabled fully automated assembly of some bacterial genomes, the assembly of human genome still produces many contigs that have to be scaffolded manually [9]. Another important downstream task is structural variant calling, where long reads can play an important role. However, present structural variant calling algorithms have low precision and recall due to noise in the reads [10]. The assembly of long segmental duplications presents another important problem where long reads can bridge repeated regions but again becomes complicated due to read errors [11].

In this paper, we propose a novel method for aligning nanopore reads that takes into account the particular structure of errors that is inherent in the nanopore sequencing process. In many of the long read aligners, the read errors are modeled using insertions, deletions and substitutions which happen at differing rates. However, in nanopore sequencing, many errors induced have structure, which is missed by viewing the errors as independent insertions, deletions and substitutions. In the nanopore sequencer, the current level depends on a *Q*-mer (a set of *Q* consecutive nucleotide bases which influence the current measurement in the nanopore). This is due to the physics of the nanopore sequencing, where a set of DNA base-pairs together influence the current output of the nanopore reader [12, 13, 14] (*e*.*g*., occupying the nanopore width). Therefore, the output current depends on a set of DNA base-pairs (*Q*-mer) influencing it. The current reading, which is used by a de-novo base caller for decoding, therefore could cause structured errors, especially between *Q*-mers that have similar outputs. This confusability between different *Q*-mers, is captured by the so-called *Q*-mer map. In Figure 1b, the median current levels for various *Q*-mers are plotted and it is clear that there is significant overlap in the current levels observed when different *Q*-mers are passed through the nanopore. These overlaps are one source of structured errors in the sequencer and can be fundamental since they can be indistinguishable by any de-novo sequencer.

**Figure 1:**
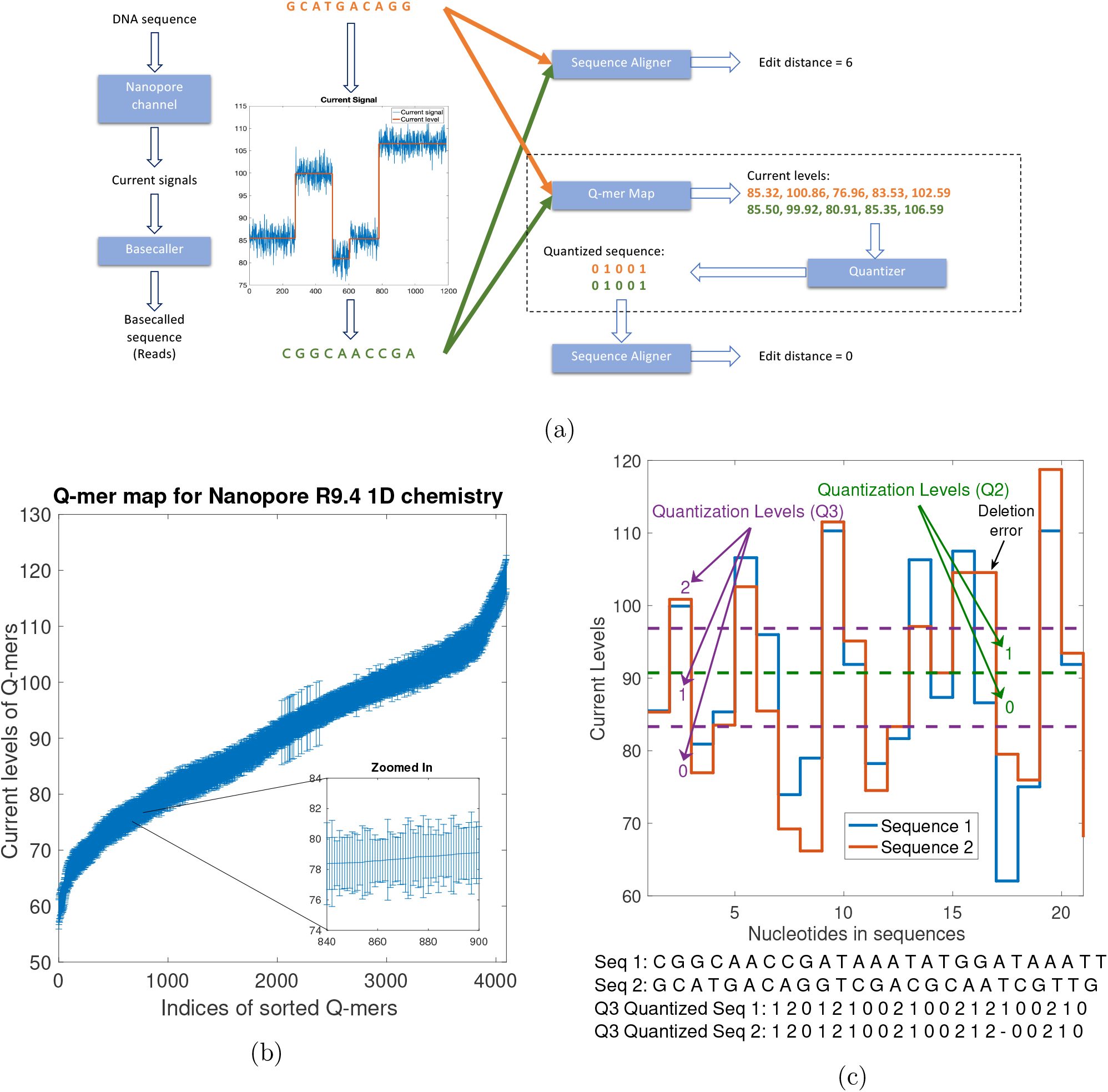
(a) An example to illustrate the error-profile in nanopore base-called reads, and the ability of QAlign to perform accurate alignment despite of the errors (the edit distance used here is to demonstrate accuracy of the alignment, however, the *nucleotide edit distance*, which is used as a metric for read-to-genome and read-to-transcriptome alignment, is computed in the nucleotide domain for the quantized alignments as well). (b) *Q*-mer map for Nanopore R9.4 1D flow cell (for *Q* = 6). It represents the physics of nanopore. The median current value along with the standard deviation (as error bars) are plotted for every 6-mers in the *Q*-mer map for R9.4 1D nanopore flow cell. Note that the difference between the median current levels of any two consecutive *Q*-mers is very small. (c) An example showing the two different nucleotide sequences have similar current levels (therefore similar quantized sequences).

The novel alignment strategy that we propose takes into account the structure of the *Q*-mer map in order to perform better alignment. In Figure 1a, we give an example where a DNA sequence (GCATGACAGG) gets wrongly sequenced as a completely different sequence (CG-GCAACCGA) due to this error mode of the nanopore sequencer. Ideally, we would like to maintain the list of “equivalent” *Q*-mers that could have plausibly caused the observed current readings. However, this is infeasible as this would entail changing the de-novo sequencing process itself to output either multiple possible reads, or give soft information about different possibilities. This is difficult, as sophisticated de-novo sequencing have been developed using artificial neural networks, which have been optimized for read error-rate performance [15]. Moreover, for a modular approach, we would not want to change the de-novo sequencer for different down-stream applications. Therefore, we take a *different* approach to resolve this problem, by using the de-novo sequenced read as the *input* to our strategy. We then deterministically *convert* this de-novo sequenced nucleotide read into a current value using the *Q*-mer median current level of the corresponding *Q*-mer (i.e. the *Q*-mer map as in Figure 1b). We further *quantize* these resulting current values from continuous values into properly chosen discrete levels. This is illustrated in Figure 1c. In this work we use 2 to 3 levels of discrete values for the quantization, which is determined based on the *Q*-mer map. Now, given this new discrete representation of the de-novo reads, we develop the new alignment algorithm, whose workflow is illustrated in Figure 1a.

A natural question is why this should help, since we are processing the de-novo reads which are erroneous, and we are *not* using any additional soft information, such as raw current values from the nanopore reads themselves. The basic insight is that the translation of the nucleotide reads to current levels enables grouping together reads that are confusable given the structure of the *Q*-mer map of the nanopore sequencer. For example, when we have two reads illustrated in Figure 1c, if the de-novo sequencer has chosen one of the two equally-likely sequences as the nucleotide read, it is clear that the alternate read, which has significant edit distance (in the nucleotide domain) is actually quite close when viewed from the lens of the *Q*-mer map, as captured by our quantized conversion process. Therefore, this process naturally groups together reads that could have been confused, and uses this as the input to our alignment algorithm, QAlign. Therefore, this reduces the effect of the errors by recognizing one structure in the error process. Note that QAlign builds an overlay layer on top of any alignment algorithm in order to align based on current levels implied by the reads instead of directly aligning the reads. Though we illustrate our ideas using the Minimap2 aligner [16], this principle can be implemented with *any* other long-read aligner such as GMAP [17].

We show that QAlign gives rise to significant performance improvements across a variety of alignment tasks including read-to-genome, read-to-read and read-to-transcriptome alignment as well as different datasets spanning from R7 nanopore sequenced data (Supplemental Figure 6) to R9.4 data.

QAlign shows significant improvement in read-to-genome alignment rates for datasets where Minimap2 alignment rate is low (improving up to around 90% for four real datasets). Furthermore, the alignments are also of higher quality: QAlign shows up to around 18% lower normalized edit distance than Minimap2 as well as longer alignments.

For read-to-read alignments, QAlign is able to align around 3.6% more overlaps between read pairs with a high overlap quality (refer to Methods for a description of the overlap quality) where Minimap2 is either unable to align the read overlaps or aligns with a low overlap quality. We show that a hybrid alignment strategy which combines QAlign and Minimap2 can improve the metric even further to around 4.6% (Supplemental Figure 14).

For read-to-transcriptome alignments, our method achieves 90% alignment rate as opposed to 82.6% with mouse 2D reads and 75.4% as opposed to 51.6% with Human 1D reads. Furthermore, the alignments are also of higher quality: QAlign shows 13.27% lower normalized edit distance than Minimap2 as well as longer alignments for Human 1D data.

In this study, we focus on the improvement of long read (in particular the Nanopore long read) alignment. To the best of our knowledge, there is no existing aligner, specifically designed to handle the error modes introduced in nanopore sequencing. There is, however, some work on incorporating the nanopore current levels in downstream tasks including post-processing of assembly by Nanopolish [18]. Nanopolish has demonstrated that utilizing the current levels can reduce assembly errors. The major difference of our work with Nanopolish is the level at which the current-level information is taken into account. Since we take into account current-level information while performing alignment, we are able to get substantially more overlaps which can lead to potentially better assembly of contigs whereas Nanopolish is only able to correct fine errors.

## 2 Methods

The QAlign strategy consists of two steps including the conversion of the nucleotide sequences to quantized (e.g. 2 levels or 3 levels) sequences in the first step. The next step is the alignment of the quantized sequences for various alignment tasks such as read-to-genome, read-to-read, and read-to-transcriptome.

### 2.1 Quantization

The nucleotide sequences are inferred from the nanopore current signals by base-callers, therefore, using a *Q*-mer map to translate the base-called sequences to the current levels implicitly maintains all of the “equivalent” base-called sequences that could be inferred from the observed current levels. These current levels can be quantized to an alphabet of finite size (Figure 1a,c).

Mathematically, the quantization process is as follows. Let Σ = {*A, C, G, T*} be the alphabet of nucleotide sequences. For a symbol *x ∈* Σ, let 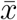 be the Watson-Crick complement of *x*. A string *s* = *x*_1_*x*_2_ …*x*_*n*_ over Σ is called a *DNA sequence*, where |*s*| = *n* is the string length and the *reverse complement* of *s* is 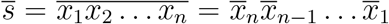. Let *p*(*s*) be a list of all *Q*-mers (e.g. *Q*=6) in the string *s*, sorted by their occurances. For example, *p*(*s*) = *k*_1_*k*_2_ … *k*_*n−Q*+1_ and each *Q*-mer *k*_*i*_ = *x*_*i*_*x*_*i*+1_ … *x*_*i*+*Q−*1_ for *i* = 1, 2, …, *n − Q* + 1. Now, we define *f* : Σ^*Q*^ → ℝ as the *Q*-mer map ^1^, which is a deterministic function that translates each *Q*-mer (*k*_*i*_) to the (median) current level (Figure 1b). Now, let *C*(*s*) = *c*_1_*c*_2_ … *c*_*n−Q*+1_ be the sequence of the current levels, so that *c*_*i*_ = *f* (*k*_*i*_) for *i* = 1, 2, …, *n − Q* + 1. The current sequence *C* can be further quantized into *w*(*s*) = *q*_1_*q*_2_ … *q*_*n−Q*+1_ by applying a thresholding function *q*_*i*_ = *g*(*c*_*i*_). The thresholding can be binary (*q*_*i*_ ∈ {0, 1}) or ternary (*q*_*i*_ ∈ {0, 1, 2}) (Figure 1c). We define 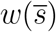 as the reverse complementary of a quantized sequence *w*(*s*), so 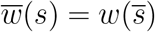.

### 2.2 Alignment

We can now use the aligners (e.g. Minimap2) to align the quantized sequences. It is important to note that these aligners inherently performs the alignment of the query sequence (e.g. *s*_1_) to the reference sequence (e.g. *s*_2_) and also aligns the *reverse complement* 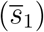 to the reference (*s*_2_). For the corresponding quantized sequences, aligners need to align the query sequence (e.g. *w*_1_) and its reverse complementary (e.g. 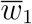) *explicitly* to the reference (e.g. *w*_2_), in order to take care of the *Q*-mer map for both *w*_1_ and 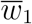 properly.

The performance of such an aligner can be evaluated by comparing the alignments of the nucleotide sequences *s*_1_ onto *s*_2_ to the alignments of their quantized sequences *w*_1_ onto *w*_2_ union with 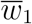 onto *w*_2_, respectively, using appropriate performance evaluation metrics.

#### 2.2.1 Read-to-Genome Alignment

We apply QAlign to the task of read-to-genome alignment. Given a nucleotide read *r* and the reference nucleotide genome *G*, we first obtain *r*^*Q*^ (the quantized *template* strand of the read, *r*^*Q*^ = *w*(*r*)) and 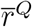 (quantized *reverse complement* strand of the read, 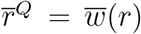 from *r*, and obtain *G*^*Q*^ (quantized reference genome) from *G*. We next align *r*^*Q*^ and 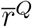 respectively to *G*^*Q*^ using Minimap2. Lastly, we aggregate the results from both the template and reverse complement outputs to determine the best alignment for each read.

Note that the quantized alignment procedure differs from the direct nucleotide alignment process in two ways. First, the nucleotide alignment does not require Minimap2 to additionally align 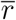 to *G* explicitly. Second, the quantized alignment employs a different seed length (e.g. minimizer length k in Minimap2) to ensure that the computation time for quantized alignment is similar as nucleotide alignment (see Supplemental Table 1 for the details of computation time versus seed lengths).

**Table 1:** Comparison for the percentage of well-aligned reads onto genome, and slope of the regression line (for normalized edit distance comparison plot of *Q*2 vs nucleotide alignments) with randomly sampled reads for each datasets. The slope of the regression line shows the average gain in the normalized edit distance.

| Dataset (No. of sampled reads) | Method of alignment | Percentage well-aligned reads | Slope of Regression line |
| --- | --- | --- | --- |
| K. Pneumoniae R9.4 1D (1k) | nucleotide | 79.4 | 0.8181 |
| | $Q2$ | 88.70 | |
| E. Coli R9.4 1D (1k) | nucleotide | 79.2 | 0.9584 |
| | $Q2$ | 84.20 | |
| E. Coli R9 2D (1k) | nucleotide | 82.6 | 0.9627 |
| | $Q2$ | 91.8 | |
| Human R9.4 1D (50k) | nucleotide | 85.70 | 0.9696 |
| | $Q2$ | 87.95 | |
| Simulated Human with Deep Simulator [23] (10k) | nucleotide | 69.04 | 0.8527 |
| | $Q2$ | 84.35 | |

We define several terms that are crucial for later performance analyis, mainly including *well-aligned, normalized edit distance*, and *normalized alignment length*.

Consider in Figure 2a, *Read 1* aligns at location *i*_1_ through *j*_1_ on the genome (we can get these locations from Minimap2 output). We say that the read is *well-aligned*, if at-least 90% of the read is aligned onto the genome (i.e., *j*_1_*−i*_1_*≥*0.9(length(*Read 1*))), and has either the *(approximate) normalized edit distance from Minimap2* (i.e. number of unmatched bases, normalized with read length, based on Minimap2 output) is less than a threshold value or the mapping quality from Minimap2 is high (greater than 20). The filtering for the well-aligned reads using this distance and mapping quality is incorporated to eliminate some spurious alignments from Minimap2. Note that the (approximate) normalized edit distance from Minimap2 is specific to nucleotide or quantized alignment. For example, for nucleotide sequences the value returned by Minimap2 is in nucleotide domain; the value returned by Minimap2 is in *Q*2 domain for the *Q*2 sequences. Therefore, different filtering threshold values are used - 0.48 for nucleotide sequence, 0.25 for *Q*2 sequence and 0.35 for *Q*3 sequence (Supplemental Figure 18 and 19).

**Figure 2:**
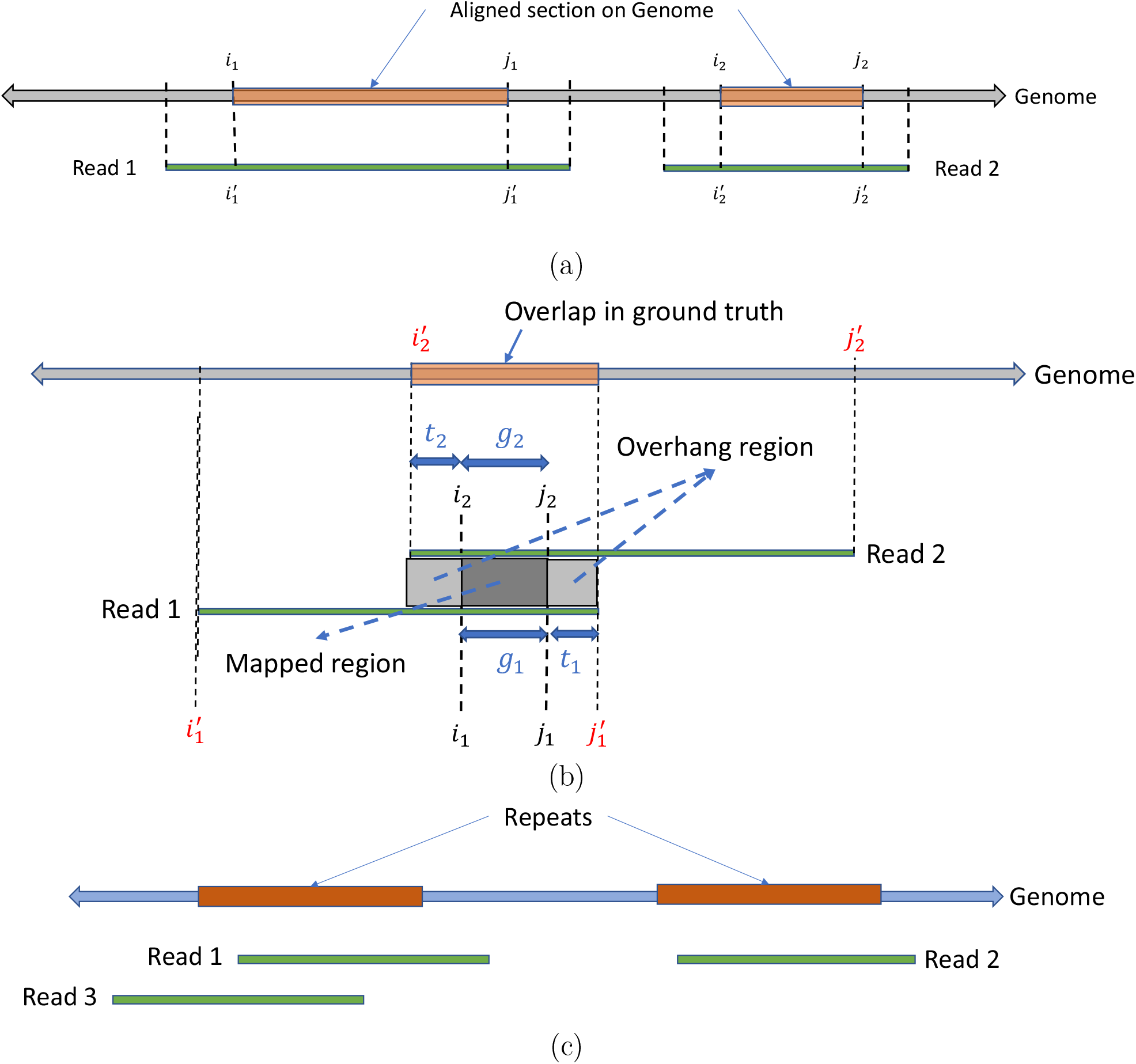
(a) An example of read-to-genome alignment. (b) An example of read-to-read alignment. (c) An example for head-to-tail alignments between reads.

In order to compare the quality of the alignments at fine-grained level, we further define *Normalized edit distance*^2^. The normalized edit distance for nucleotide alignment is 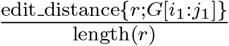 and for quantized alignment is 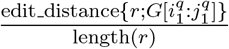, where *i*_1_, *j*_2_ and 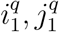 are locations obtained from nucleotide and quantized alignment respectively. Similarly, we define *Normalized edit distance of aligned read* for nucleotide alignment as 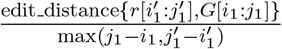 (Figure 2a) and for the quantized alignment as 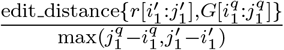. As evident from the definitions, these metrics for both nucleotide and quantized alignment are calculated all in nucleotide domain (unlike the approximate normalized edit distance from Minimap2, which is domain specific). Specifically, for quantized alignment, we leverage the information about the alignment location on genome (i.e. 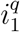 and 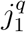) to calculate the normalized edit distance between the nucleotide read and the corresponding aligned section on the nucleotide genome.

Another metric at fine-grained level is *normalized alignment length*, which is the ratio of the length of the section on genome where a read aligns to the length of the read. It is 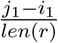 for nucleotide alignment, and 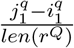 for quantized alignment. A contiguous alignment tends to have this metric as 1.

We have been discussing the nanopore 1D reads for read-to-genome alignment so far. There are also 2D reads (e.g. *r*), which are the consensus calling using the 1D reads from both the *template* strand (e.g. *r*_*t*_) and the *complement* strand (e.g. *r*_*c*_). For the read-to-genome alignment algorithm of the 2D reads using QAlign, the experiment pipeline has been modified so that the error profile introduced in the sequencing of the 1D reads can be mitigated. Specifically, the quantized reads from both the template strand (e.g. 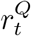) and the complement strand (e.g. 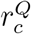) are aligned to the quantized genome (e.g. *G*^*Q*^) individually using Minimap2. The union of the two alignments^3^ is considered as the output of the QAlign algorithm. In case there is no overlap in the alignments of the reads from the individual strands, both the genome sections are given as the output of the QAlign algorithm, as the 2D consensus read might align to either of these sections. Since QAlign yields the genome section as the union of the two alignments, it could be much larger (nearly twice) than the read length. Therefore, the genome section needs to be further refined by the local alignment of the 2D consensus read onto the section. The performance evaluation of QAlign is mainly based on the normalized edit distance between the 2D consensus read and the refined genome section (the results using this method for the 2D read alignment onto genome are discussed in Supplemental Figure 3 and 4).

**Figure 3:**
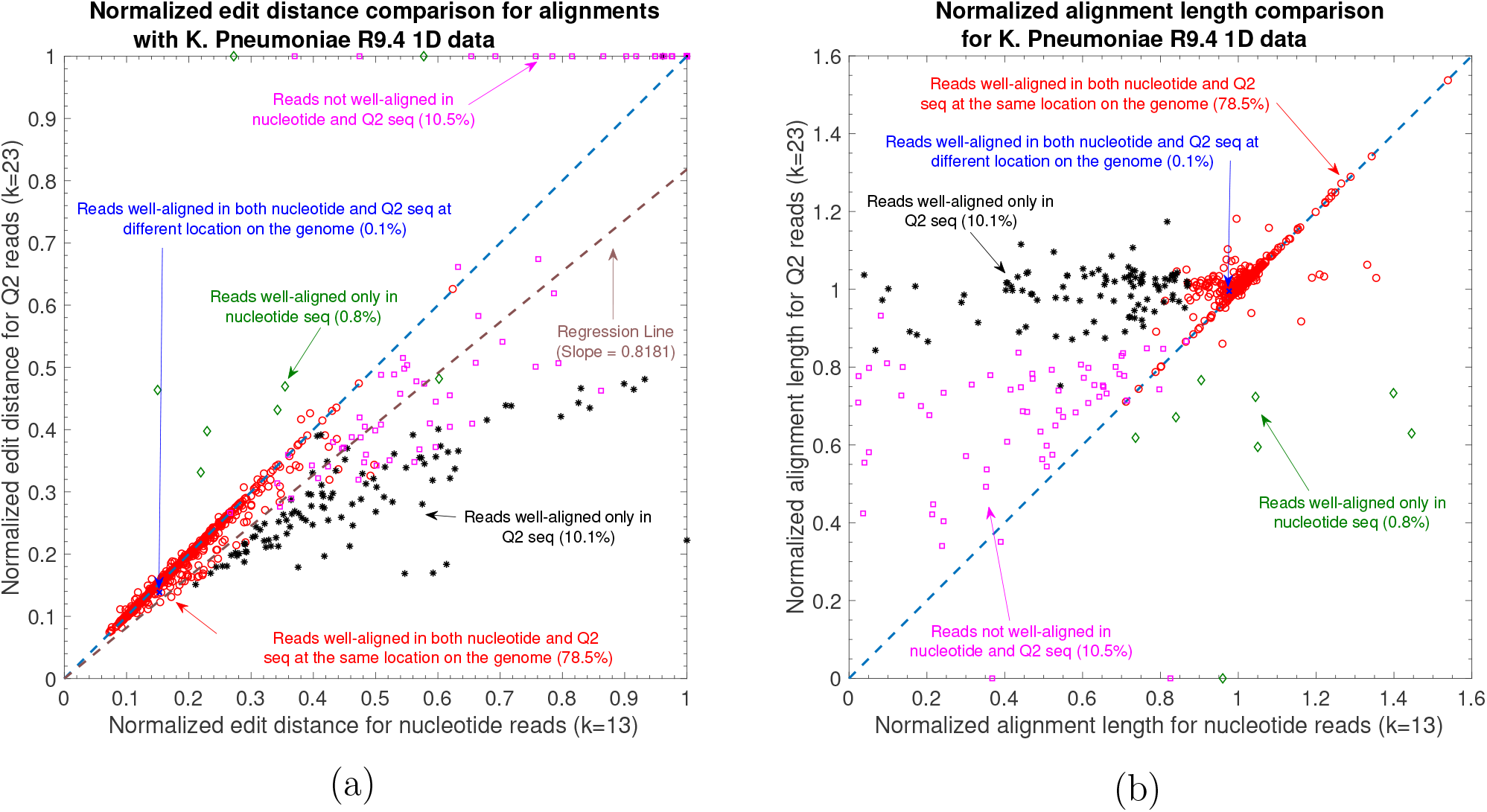
Nanopore long DNA reads alignment onto Genome. (a) Comparison of normalized edit distance for K. Pneumoniae R9.4 1D reads data. Smaller values for *normalized edit distance* is desirable as it represents better alignment. The slope of the regression line is *<* 1, therefore, representing better alignments with *Q*2 than nucleotide alignments for same reads on average. (b) Comparison of normalized align-length on genome for K. Pneumoniae R9.4 1D reads data. Normalized alignment length of 1 is desirable as it represents that entire read is aligned. Majority of the reads are above *y* = *x* line representing longer alignment length in *Q*2 than nucleotide alignment.

**Figure 4:**
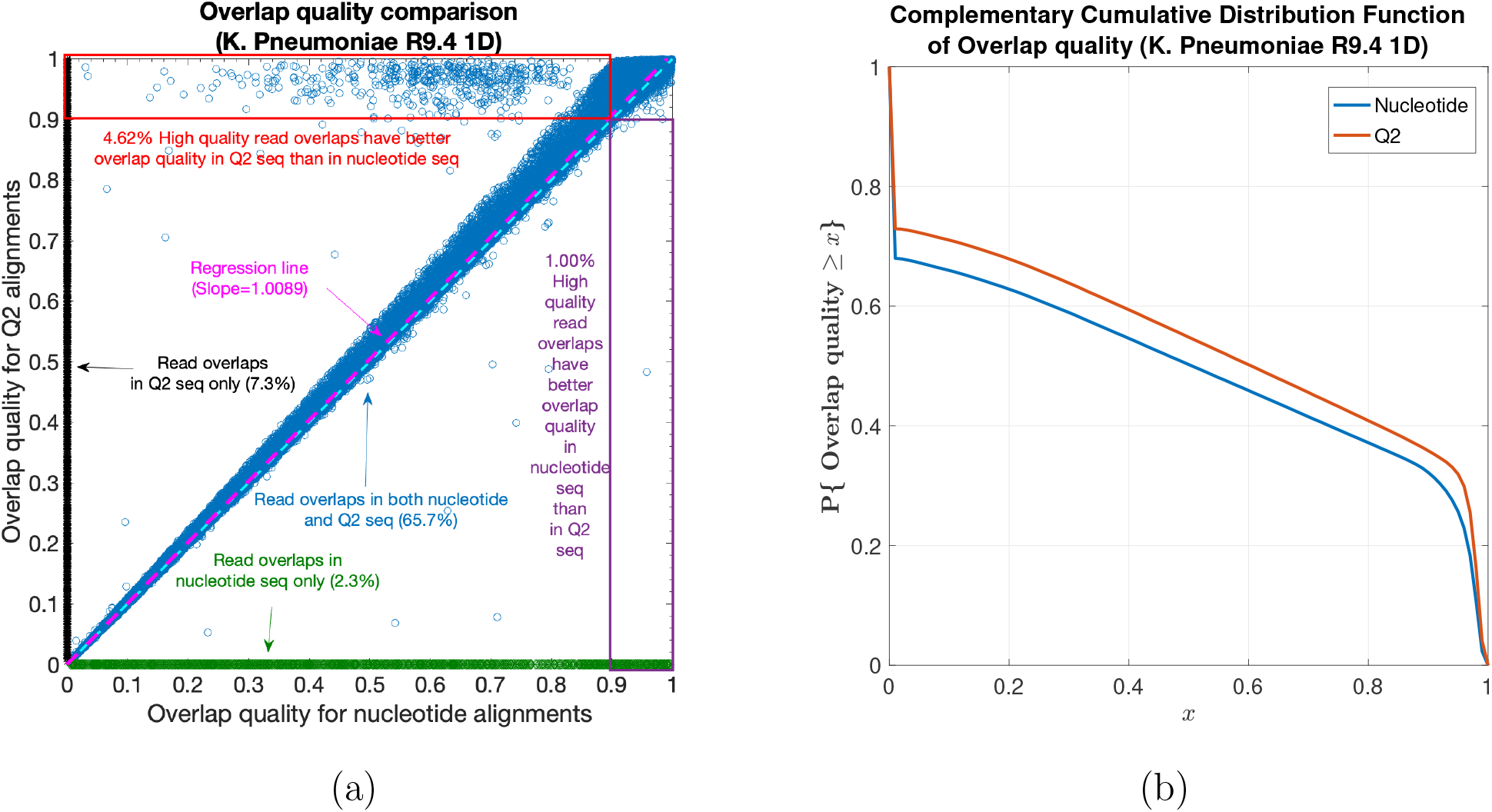
Nanopore long DNA read-to-read alignment. (a) Comparison of overlap quality for K. Pneumoniae R9.4 1D reads dataset (*Q*2 vs nucleotide). Overlap quality of 1 is desirable as it represents the alignment of the algorithm matched the alignment in the ground truth exactly. Therefore, slope of the regression line *>* 1 represents better overlap quality of *Q*2 alignments than nucleotide alignments on average. (b) Complementary CDF of overlap quality for K. Pneumoniae R9.4 1D reads dataset. *Q*2 curve is strictly above the curve for nucleotide, therefore, demonstrating better overlap quality for *Q*2. Area under the curve gives an average overlap quality which is higher for *Q*2.

#### 2.2.2 Read-to-Read Alignment

We apply QAlign to read-to-read alignment as the second alignment task, which provides overlaps between reads that are typically necessary for genome assembly.

The alignments between the nucleotide reads *r*_1_ and *r*_2_ (or between the quantized reads 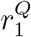 and 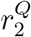) are obtained using Minimap2 as well. Similar to read-to-genome alignment, the quantized alignment not only aligns 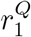 to 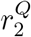, but also needs to align 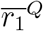 to 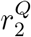. In addition, the quantized alignment employs a seed length (e.g. the minimizer length ‘`k`’ in Minimap2) different from nucleotide alignment so that the computation time in both nucleotide and quantized regimes is maintained to be similar (see Supplemental Table 2 for detailed analysis of computation time versus `k`).

**Table 2:** Comparison for precision, recall and and average overlap quality for read-to-read alignment for four different datasets. Average overlap quality is computed as the area under the complementary CDF curve of overlap quality.

| Dataset | Method of alignment | Precision (%) | Recall (%) | Avg. Overlap quality |
| --- | --- | --- | --- | --- |
| K. Pneumoniae R9.4 1D | nucleotide | 97.47 | 67.99 | 0.4908 |
| | $Q2$ | 96.93 | 72.92 | 0.5360 |
| | $Q3$ | 97.49 | 69.52 | 0.5053 |
| E. Coli R9.4 1D | nucleotide | 99.20 | 62.27 | 0.4688 |
| | $Q2$ | 99.06 | 62.60 | 0.4803 |
| | $Q3$ | 99.23 | 63.87 | 0.4811 |
| E. Coli R9 2D | nucleotide | 98.94 | 59.42 | 0.5339 |
| | $Q2$ | 96.97 | 65.47 | 0.5914 |
| | $Q3$ | 98.99 | 62.46 | 0.5615 |
| Simulated Human with Deep Simulator [23] | nucleotide | 75.72 | 41.91 | 0.3888 |
| | $Q2$ | 75.34 | 53.10 | 0.5100 |
| | $Q3$ | 76.08 | 54.19 | 0.5174 |

For the algorithm evaluation purpose, we need to have the ground truth, which is unknown. One way to judge the quality is to compute the normalized edit distances of alignment overlaps. However, this is not only computationally expensive but also suffers from false alignments between reads from repeated regions. Instead, we leverage the read-to-genome alignments to build the ground truth for read-to-read alignment. Specifically, all of the reads are firstly aligned to the genome via both the nucleotide alignment and the quantized alignment. The reason behind performing the read-to-genome alignment in both the nucleotide and the quantized domain is to ensure that more read alignments are captured, as there can be some alignments that are captured/contiguous only in quantized alignments and vice-versa. For the experiments, we focus on a section of the genome *G* (say, *G*_1_=*G*[1:1000000]) to find all the reads aligning onto *G*_1_. Assume there are *n*_1_ and *n*_2_ number of reads aligned to *G*_1_ in nucleotide domain and in quantized domain, respectively, with normalized edit distance (in nucleotide domain for both methods) less than 0.48, which indicates the found alignment of the reads are better than the alignment of two random DNA sequences (see Supplemental Figure 18). Now, we randomly choose *n* reads from a union of *n*_1_ and *n*_2_ reads such that 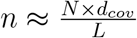, where *d*_*cov*_ is the required coverage depth, *N* is the length of the genome section *G*_1_ (i.e. 1000000), and *L* is the average length of the *n*_1_ ∪ *n*_2_ reads.

To estimate the ground truth, consider the alignment of *Read 1* (*r*_1_) and *Read 2* (*r*_2_) onto the genome as shown in Figure 2b, where the alignment locations of the reads on the genome are 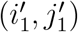 and 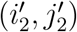, respectively. We say that the reads are *overlapping in the ground truth* if there is an overlap (denoted as *l*) of at least 100 bases, where 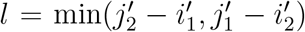. For reads that have overlaps in both nucleotide and quantized alignment, denoted as (*l*^*nucleotide*^) and (*l*^*Q*^) respectively, the larger one is chosen as *l* = max(*l*^*nucleotide*^, *l*^*Q*^)).

Provided the ground truth, we can compute the Precision and Recall to make a comparison between the two methods. Precision is defined as the fraction of overlaps in the ground truth among the overlaps determined by the algorithm. Recall (also known as *sensitivity*) is the fraction of overlaps in the ground truth that are determined by the algorithm.

The read-to-read alignment will label two reads to have an overlap (different from the overlap used to find ground truth) if the length of the ‘Mapped Region’ is at least 90% of the ‘Mapped Region’ plus the ‘Overhang Region’ (Figure 2b, i.e., *g*_1_ *≥* 0.9(*g*_1_+*t*_1_+*t*_2_) and *g*_2_ *≥* 0.9(*g*_2_+*t*_1_+*t*_2_), or equivalently *t*_1_ + *t*_2_ *≤* 0.1(min(*g*_1_, *g*_2_)). For evaluation, we define another metric called the *overlap quality* (denoted as *X*) as 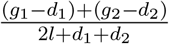 where ^4^ the overlap quality measures how well the reads are aligned with respect to each other, compared to the alignment in ground truth. Ideally, it is desired to have the value of overlap quality close to 1. We also define the *average overlap quality*, which is the expected value of the overlap quality (i.e. 𝔼[*X*] = ∫ℙ{*X* > *x*}*dx*), or the area under the complementary CDF of *X*.

It is possible that two reads will be falsely aligned especially when they are from repetitive regions. To remedy this, we only consider head-to-tail alignment between reads. For example in Figure 2c, three reads *Read 1, Read2* and *Read 3* have been sequenced where *Read 1* and *Read 2* are from repetitive regions. Consequently after read-to-read alignment, there will be an overlap between *Read 1* and *Read 2* that can be filtered out since it is not a head-to-tail alignment. However, there will also be another false positive overlap between *Read 3* and *Read 2*, which will not be removed as it satisfies the head-to-tail alignment condition. In order to further reduce the number of false positives of read-to-read alignments, the (approximate) normalized edit distance provided by Minimap2 is used for extra filtering (see Supplemental Figure 18 and 19).

In addition to reduce false positives, the read-to-read alignment results can be further improved by implementing an Ensemble model, which is able to capture the best alignment (e.g. longer length of ‘Mapped Region’) from both methods of the nucleotide alignment and the quantized alignment, as well as to incorporate the alignments that are complementary in either method (see Supplemental Figure 14).

#### 2.2.3 Read-to-Transcriptome Alignment

Applying QAlign strategy to the third task of the RNA read-to-transcriptome alignment is analogous to the DNA read-to-genome alignment. This is not the spliced alignment of the reads to the genome; instead all of the RNA reads are aligned to the transcriptome. Since the ground truth is unknown for the alignments, we use *normalized edit distance*, and *normalized alignment length* as the evaluation metric.

## 3 Results

In this section, we demonstrate and discuss the results for (i) DNA Read-to-Genome alignment, (ii) DNA Read-to-Read alignment, and (iii) RNA Read-to-Transcriptome alignment using QAlign and Minimap2.

### 3.1 Datasets

#### Datasets for DNA-seq alignments

We use five datasets for DNA read-to-genome and read-to-read alignment: (1) MinION sequencing of K. Pneumoniae DNA using R9.4 1D flow cell [19], (2) MinION sequencing of E. Coli DNA using R9 2D flow cell [20], (3) MinION sequencing of E. Coli DNA using R9.4 1D flow cell [21], (4) MinION sequencing of Human genome using R9.4.1 flow cell [22], and (5) Simulated read data from GRCh38 chromosome 1 using Deep Simulator [23] for benchmarking the performance of QAlign.

#### Datasets for RNA-seq alignments

The experiments are based on the RNA reads obtained by MinION sequencing of human cDNA using R9.4 1D flow cell [24], and MinION sequencing of mouse cDNA using R9.4 2D flow cell [25] (SRA access No. SRR5286961).

Compared to DNA-seq datasets, RNA-seq datasets carried out using nanopore sequencing are relatively rare. We select these datasets because they have relatively complete reference transcriptome (e.g. for human there are 200,401 annotated transcripts [26] and for mouse 46,415 annotated transcripts from UCSC Genome Browser [27]), and the corresponding *Q*-mer map models [28] are available for quantization.

### 3.2 DNA Read-to-Genome Alignment

The alignment of DNA reads to the genome is a task with wide-ranging applications in sequencing experiments. It is a required step in variant calling pipelines [29], in particular structural variant calling can benefit significantly from long reads offered by the nanopore sequencing platform [10]. It is also useful in calling variants in long segmental duplications [11], where long duplications necessitate long reads to resolve the repeat ambiguity. Another application for DNA read-to-genome alignment appears in reference matching - for example, in meta-genomics, in estimating which reference species is present in the sample.

The results are illustrated in the Figure 3 and Table 1. At a coarse level, the performance is measured by the fraction of the reads that have been well-aligned by the algorithm. A read is said to be well-aligned if at-least 90% of the read is aligned to genome and has either the (approximate) normalized edit distance from Minimap2 (i.e. number of unmatched bases, normalized with the read length) below a threshold value or the mapping quality from Minimap2 is high (see Methods). QAlign is shown to significantly improve the fraction of well-aligned reads - in particular, in the K.Pneumoniae R9.4 1D dataset, this metric improves to 88.7% from 79.4%. In the E. Coli R9.4 1D dataset, it improves to 84.2% from 79.2%; in the E. Coli R9 2D dataset, the numerics improves to 91.8% from 82.6%, and for the human R9.4.1 dataset, it improves to 87.95% from 85.70%. For the benchmark with the simulated data, the numerics improves to 84.35% from 69.04% (refer to Table 1).

The results in Figure 3a-b compares the quality of the alignments using Minimap2 and QAlign at a fine-grained level for the K. Pneumoniae dataset (plots for other datasets are available in Supplemental Figure 1-10).

Specifically, Figure 3a compares the normalized edit distance for QAlign and Minimap2. The normalized edit distance is the edit distance between the entire read and the aligned section on the genome normalized with the length of the read, in nucleotide domain for *both* nucleotide alignment and quantized alignment (Q2). In case of Q2, the information of the location of the alignment on the genome is leveraged from the alignment between the quantized read and the quantized genome first, and the edit distance is computed between the corresponding nucleotide read and the aligned section on the nucleotide genome (see Methods for details). Intuitively, the normalized edit distance gives a measure of how close the two sequences are. Therefore, the smaller the normalized edit distance, better is the alignment. In addition, the *normalized edit distance* for the reads that have *normalized edit distance of aligned reads* more than 0.48 is set to 1 (We noticed that the normalized edit distance between a pair of random DNA sequences is above 0.48, refer to Supplemental Figure 18). Therefore, the figure represents only those alignments that are better than alignment of random DNA sequences.

To better visualize the results, we group alignments with different colors and marks for different conditions. The red circles in Figure 3a-b represent the reads that are well-aligned in both nucleotide and *Q*2 alignments and at nearly the same location on the genome. The blue cross represent well-aligned reads in both *Q*2 and nucleotide alignments but at different location on the genome or on a different chromosome. The black asterisks are the reads that are well-aligned in *Q*2 only, *i*.*e*., in nucleotide alignments, the alignment of these reads are either missing or does not satisfy the definition of the well-aligned reads. The green diamonds are the reads that are well-aligned in nucleotide alignments only. The pink square points are the reads that are not well-aligned in both *Q*2 and nucleotide alignments. For each read, there could be multiple alignments on the genome because of the repeats in the genome, but we consider the alignment that has the minimum edit distance amongst all of them for the evaluation in these plots.

Figure 3a shows that the normalized edit distance is overall smaller for *Q*2 alignments than nucleotide alignments. The better alignment in *Q*2 is also evident from the slope of the regression line in Figure 3a. It shows that on average *Q*2 alignments has 18.19% improvement in terms of the normalized edit distance than the nucleotide alignments.

The results for another fine-grained metric are shown in Figure 3b, which compares the normalized alignment length on genome in *Q*2 to the normalized alignment length on genome in nucleotide alignments. The normalized alignment length is the ratio of the length of the section on genome where a read aligns to the length of the read. There are 10.1% reads that are well-aligned in *Q*2 only (the black asterisks), and the normalized alignment length is close to 1 in *Q*2 but it is much less than 1 in nucleotide alignments, therefore representing several non-contiguous alignments in nucleotide domain that are captured in *Q*2. The normalized edit distance for such reads in *Q*2 is much less than the normalized edit distance for the same reads in nucleotide alignments. Similar results are observed across different datasets as evident from the slope of the regression line for normalized edit distance comparison between *Q*2 and nucleotide alignments shown in Table 1.

### 3.3 Read-to-Read alignment

Alignment of genomic reads to other reads is a basic primitive useful in many settings. For example, this is a first step in many overlap-layout-consensus (OLC) assemblers [1]. A key challenge in read-to-read assembly is the increased error rate that the aligner has to deal with. For example, if two reads *R*_1_, *R*_2_ are sampled from the same region of the genome, each may be within 15% edit-distance of the reference genome (assuming a 15% error-rate), however, the edit distance between *R*_1_ and *R*_2_ can be up to 30% leading to an effective doubling of the error-rate. Long-reads hold the promise of fully-automated assembly but is currently feasible only when for bacterial genomes [30]. For complex mammalian genomes, long repeats fragment assembly [31] and more accurate alignment can help alleviate this problem.

The results for read-to-read alignment are illustrated in the Figure 4a-b, and Table 2. Table 2 summarizes the precision, recall, and average overlap quality for different methods used (namely, nucleotide, *Q*2, and *Q*3) to find the alignments between the overlapping reads across different datasets. It is evident from the table that *Q*2 provides higher recall and average overlap quality than nucleotide alignments at the cost of a bit lower precision. *Q*3, on the other hand, shows better recall and average overlap quality than nucleotide alignments at similar precision.

For a fine-grained evaluation, Figure 4a shows overlap quality comparison for the quantized (*Q*2) alignments versus nucleotide alignments using the K. Pneumoniae dataset. The blue circles in the figure represent the overlaps that are aligned ^5^ in both QAlign and Minimap2. The black asterisks (along the line *x*=0) represent the overlaps that are aligned only in *Q*2 and not aligned in nucleotide, whereas the green diamonds (along the line *y*=0) represent the overlaps that are aligned only in nucleotide and not aligned in *Q*2. In Figure 4a, the read overlaps that are aligned only in *Q*2 is 7.3%, whereas the read overlaps that are aligned only in nucleotide is 2.3%. Therefore, QAlign demonstrates a net gain of 5.0% in terms of the number of reads aligned by the algorithm. For the read overlaps that are aligned in both *Q*2 and nucleotide, 4.62% of the read overlaps have overlap quality more than 0.9 in QAlign but not in Minimap2 whereas the opposite holds true in only 1.0% of the read overlaps. Thus QAlign gives a net performance improvement of 3.62% over Minimap2. In addition to that, the slope of the regression line in the figure is 1.0089, therefore also illustrating better overlap quality with QAlign than Minimap2.

Figure 4b shows the fraction of reads which have overlap quality greater than *x* for the two aligners - the performance gain is seen to hold across a wide range of threshold values *x*. The area under the curve (which equals the average overlap quality) is computed for nucleotide, *Q*2, and *Q*3 alignments across all the datasets and is demonstrated in Table 2. The gain in the average overlap quality is observed using QAlign across all the datasets as evident from Figure 4b. Specifically, there is a gain of 9.2% in K. Pneumoniae dataset, when we compute it as the ratio of the average overlap quality of *Q*2 to average overlap quality of nucleotide alignments. Similarly, there is a gain of 2.5%, 10.8%, and 31.2% in the average overlap quality for the E. Coli R9.4 1D, E. Coli R9 2D dataset, and simulated human dataset, respectively.

### 3.4 Read-to-transcriptome alignment

RNA-seq is a popular sequencing technology with emerging applications including single-cell RNA-seq [32]. While short high-throughput reads may suffice to assess rough gene expression estimates, isoform level analysis is better facilitated by long nanopore reads that can straddle several exons simultaneously [7]. Here we perform the alignment of cDNA reads (complementary DNA reads extracted from reverse transcription of RNA) onto a reference transcriptome.

The results for read-to-transcriptome alignment are illustrated in Figure 5a-b, and Table 3. At a coarse level, QAlign improves the fraction of the well-aligned reads significantly. For the Human R9.4 1D dataset, the metric improves to 75.40% from 51.60%, and for the Mouse R9.4 2D dataset, it improves to 90.00% from 82.60%, as shown in Table 3.

**Table 3:** Comparison for the percentage of well-aligned reads onto transcriptome, and slope of the regression line (for normalized edit distance comparison plot for *Q*2 vs nucleotide) for two different dataset for randomly sampled reads for each dataset.

| Dataset (No. of sampled reads) | Method of alignment | Percentage well-aligned reads | Slope of the Regression line |
| --- | --- | --- | --- |
| Human R9.4 1D (2k) | nucleotide | 51.60 | 0.9125 |
|  | *Q2* | 75.40 |  |
| Mouse R9.4 2D (2k) | nucleotide | 82.60 | 0.8455 |
|  | *Q2* | 90.00 |  |

**Figure 5:**
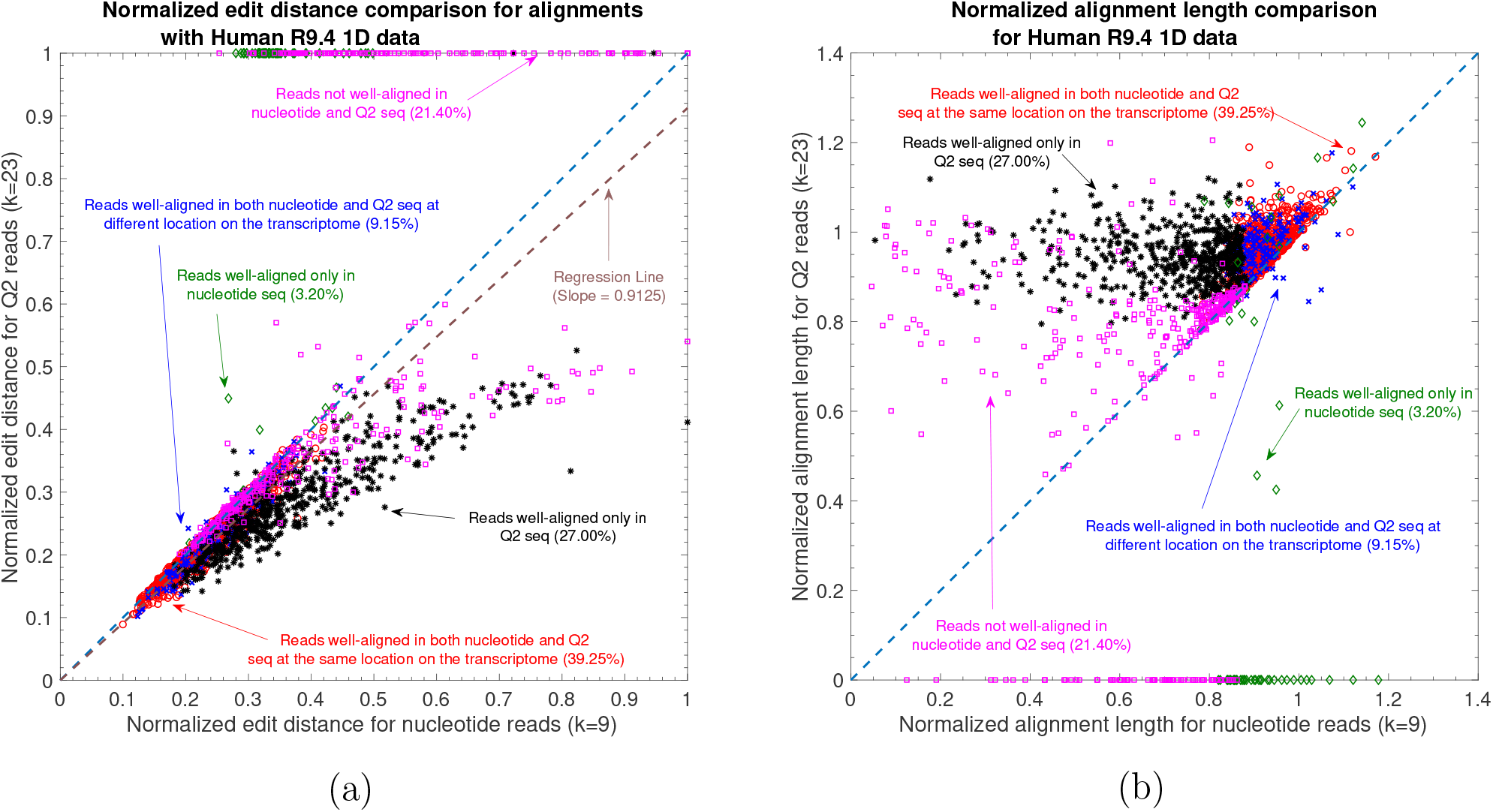
Nanopore long RNA read to transcriptome alignment. (a) Comparison of normalized edit distance for Human R9.4 1D dataset. A small *normalized edit distance* is desirable as it represents better alignment. The slope of the regression line is *<* 1, therefore, representing better alignments with *Q*2 than nucleotide alignments for same reads. (b) Comparison of normalized alignment length of the aligned sections on the transcriptome for Human R9.4 1D dataset. Normalized alignment length of 1 is desirable as it represents that entire read is aligned. Majority of the reads are above *y* = *x* line, representing longer alignment length in *Q*2 than nucleotide alignment.

At a fine-grained level, Figure 5a compares the normalized edit distance for Human R9.4 1D dataset. Note that the *normalized edit distance* is set to 1 for the reads that have *normalized edit distance of aligned reads* greater than 0.48. Therefore, the figure represents the alignments that are not “equivalent” to the alignment of random nucleotide sequences. This figure clearly demonstrates the gain of quantized alignment. Specifically, *Q*2 is able to align 27.00% more reads with 8.75% better quality than nucleotide alignments (from the slope of the regression line; a similar trend of slope of regression line using Mouse R9.4 2D dataset is shown in Table 3). In Figure 5b, the lengths of aligned chunks are compared between nucleotide and *Q*2 domain. Most of the reads gets larger aligned chunks using *Q*2 quantization. Moreover, we observe a similar trend in the alignment using the Mouse R9.4 2D dataset as shown by the slope of the regression line in Table 3.

## 4 Discussion

QAlign is a pre-processor that can be used with any long read aligner for a nanopore sequencer. It can be used for aligning reads onto genome or as a long-read overlapper or for aligning RNA-seq reads onto transcriptome. QAlign provides alignments that outperforms other aligners that uses nucleotide sequences in terms of the accuracy of the alignment at the cost of a similar computation time.

The reason for this performance improvement is because it takes into account the underlying physics of the nanopore sequencer through its *Q*-mer mapping, which could be the pre-dominant cause of the error behavior in nanopore sequencing. We demonstrated how the structure of the *Q*-mer map can be used even with only nucleotide read outputs, and without access to the current-level output of the sequencer. In particular, QAlign converts the nucleotide reads to quantized current levels (of finite alphabet size) which are then aligned using any state-of-the-art aligner. This improvement in the alignment of the long nanopore reads can be useful in several downstream applications such as structural variant calling, assembly - where the QAlign can benefit in the discovery of SVs and read overlaps that are difficult to capture because of the high error rate of nanopore reads.

The current limitation of QAlign is that it works well when we have long contiguity in the alignments. Therefore it does not perform as well in doing the spliced alignments of the RNA-seq reads onto genome while maintaining a similar computation time cost (as shown using empirical results in Supplemental Figure 20). Part of ongoing extensions is to build a deep hybrid aligner which brings together the advantages of the nucleotide alignments and QAlign.

## Author’s contributions

DJ, SK, and SD conceived the original idea and developed the project. DJ led the development of the software tool, and SM assisted in its open-source development. DJ and SM performed the analysis on the various datasets. All the authors wrote the manuscript.

## Acknowledgements

SD and DJ were supported in part by NSF grant 1705077. SM and SK were supported in part by NIH grant 1R01HG008164 and NSF grants 1651236 and 1703403.

## Footnotes

1 *Q*-mer map is determined by the chemistry of the nanopore flow cell, and is therefore dataset dependent, *i*.*e*., the *Q*-mer map for sequencing using R9 flow cell is different from *Q*-mer map for sequencing using R9.4.1 flow cell. The *Q*-mer maps used in this work are generated by [18].

2 This is different from *approximate normalized edit distance from Minimap2* to filter for *well-aligned* reads.,

3 For example, alignment regions are [0, 2] and [1, 3], and union is [0, 3].

4 Empirically *d*_1_ and *d*_2_ tend to be simply zero

5 An overlap between a pair of reads is said to be aligned by the algorithm if the Mapped region by the algorithm is at least 90% of the Mapped region plus the Overhang region (refer to the Methods section for more details).

