## Supplementary file for "QAlign: Aligning nanopore reads accurately using current-level modeling"

### Supplementary Material for "QAlign: Aligning nanopore reads accurately using current-level modeling"

#### 1 Read-to-Genome Alignment

*Methodology and performance metrics:* The ability of QAlign to align DNA reads to a reference genome is discussed here. In each experiment, we take the reads and align them to the genome both using QAlign and Minimap2. There are two different types of performance metrics: (1) Coarse performance is measured by the fraction of reads that have been *well-aligned* by the algorithm. A read is said to be well-aligned if at least 90% of the read is aligned to the genome and the *(approximate) normalized edit distance* from Minimap2 is below a threshold value (threshold values are 0.48 for nucleotide alignments, 0.25 for $Q2$, and 0.35 for $Q3$ - these values are chosen based on the empirical results shown in Supplemental Figure 18 and 19) or the mapping quality is high (greater than 20). (2) Fine-grained performance is measured in our experiments by two metrics: the *normalized alignment length* and the *normalized edit distance* between the read and the genome region that the read aligns to. We note that we do not use the particular alignment returned by the different aligners since this is not directly comparable. The normalized-edit distance is defined as the ratio of the edit distance between the genomic region aligned by the read and the entire read to the length of the read - thus, normalized edit distance of 0 corresponds to perfect alignment, and an upper bound is 1 if the read does

†DJ and SD are at the University of California, Los Angeles. SM and SK are at the University of Washington, Seattle

not return an alignment. The normalized edit distance is computed in the nucleotide domain for both the nucleotide and quantized alignments. The information about the start and the end location on the genome is leveraged to compute the edit distance in the nucleotide domain for the quantized alignments. We note that the normalized edit distance between a random sequence of genome and a given read of the same length is 0.48 (Refer to Supplemental Figure 18 for the details on the normalized edit distance between random DNA sequences).

The Minimap2 command for the read to reference alignment with nucleotide sequence is:

```
minimap2 -c -k 9 reference.fasta reads.fasta
```

The Minimap2 command for the alignment with the quantized sequences is:

```
minimap2 -c -k 23 --for-only reference_q2.fasta reads_q2.fasta
minimap2 -c -k 23 --for-only reference_q2.fasta rc_reads_q2.fasta
```

The two separate commands for the alignment with the quantized sequences are for the alignment of the quantized *template* strand of the reads to the quantized genome and the alignment of the quantized *reverse complement* strand of the reads to the quantized reference genome, respectively. We can then aggregate the results from both the outputs to determine the best alignment for each read.

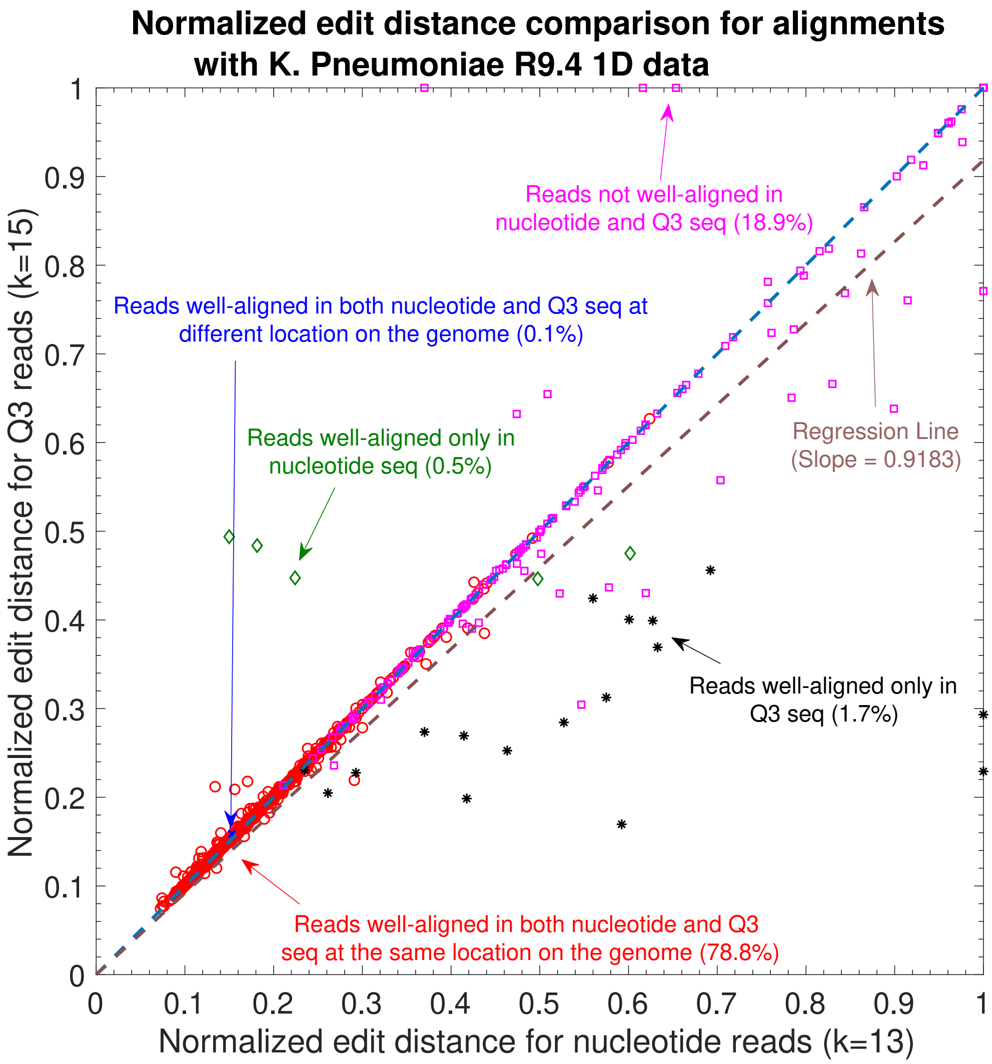


Figure 1: Normalized Edit distance comaprison for nucleotide vs $Q3$ for K. Pneumoniae R9.4 1D reads. The slope of regression line is 0.9183, indicating 8.17% reads have a smaller distance in Q3 than in nucleotide. 1.7% reads are well aligned to genome in $Q3$ only, whereas there are 0.5% reads that are well aligned in nucleotide domain only.

*Read-to-genome alignment for the 2D reads:* An alternate pipeline for the alignment of the 2D consensus reads is using the corresponding 1D reads from both the template and the complement strand and estimate the alignment location of the 2D consensus read onto the genome using the QAlign algorithm. This can be done by exploiting the nanopore physics with the 1D reads from each strands. The experiment pipeline is explained in detailed in the method section of the main paper. The results in Supplemental Figure 2 illustrates that the alignment length on the genome, which is the output of the QAlign algorithm is atleast twice the length aligned by the local alignment algorithm for the same read at no additional cost of the normalized edit distance (as shown in Figure 4 and 5 in the Supplementary paper).

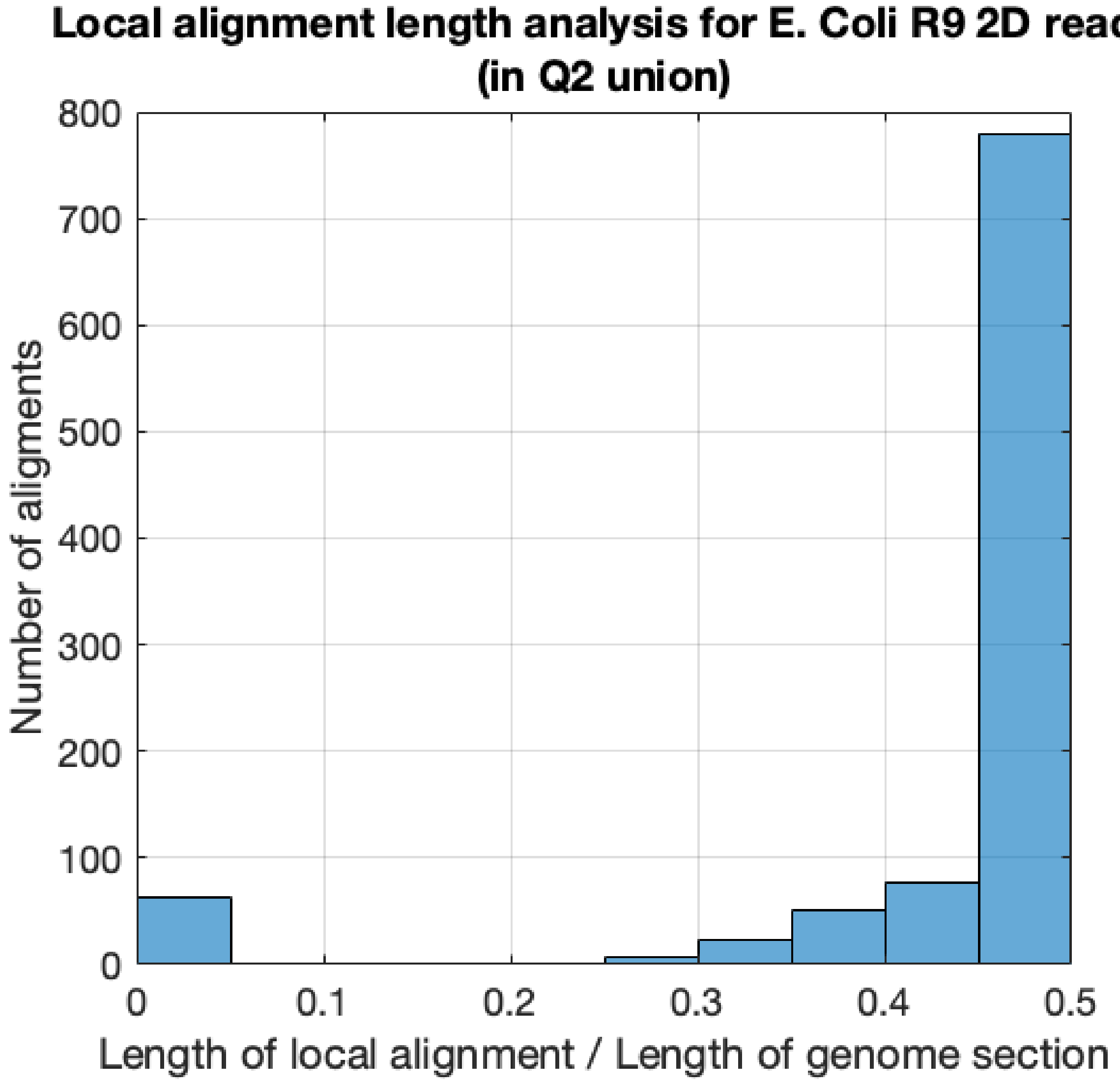


Figure 2: Hybrid approach for the 2D reads using the alignments from the 1D reads of each strand of the DNA. Length of the genome section is the output of QAlign using the alternate approach for 2D reads and it is the union of the region where the 1D reads from both the template and the complement strand aligns to on the genome. Length of local alignment is the region on this genome section where the 2D consensus reads aligns to (using a local alignment algorithm). Therefore, the output of QAlign algorithm is atleast twice the length of the local alignment.

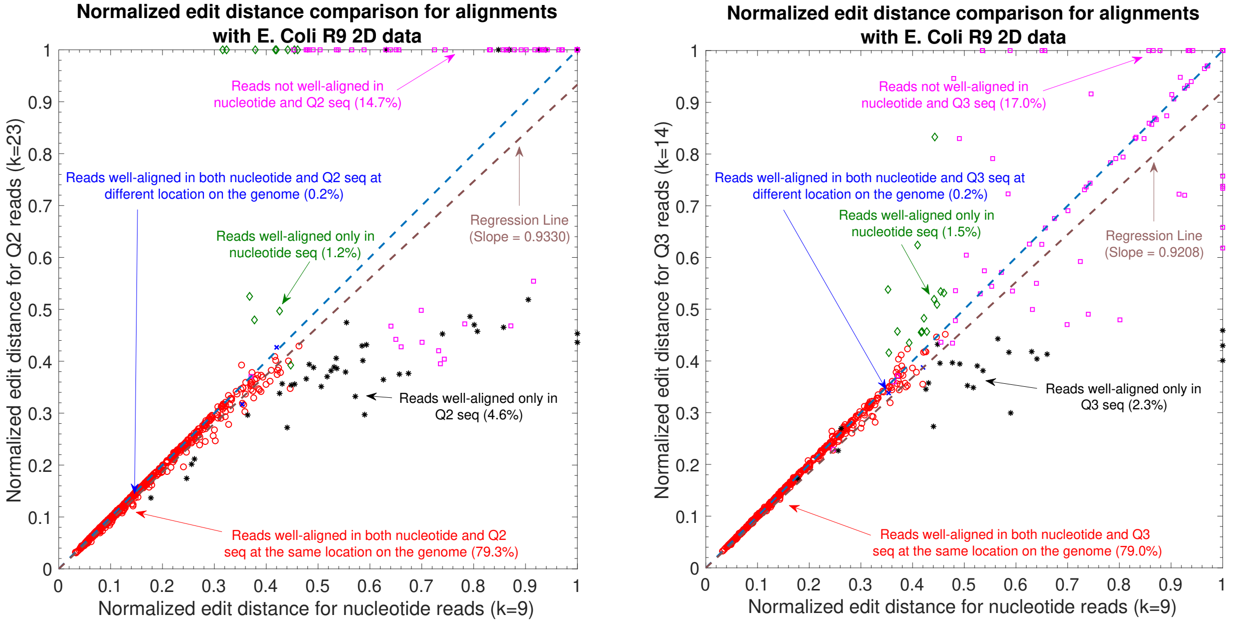


(a) (b)

Figure 3: The alignment of long nanopore DNA-Seq R9 2D reads onto E. Coli reference genome using the same pipeline for 2D reads, i.e., the 2D reads are translated to the quantized current sequences without any knowledge of the 1D reads. (a) The plot shows the normalized edit distance comparison for nucleotide vs $Q2$ in the 2D reads experiment setting. The slope of the regression line is 0.9330, indicating 6.7% well aligned reads have a better normalized edit distance in $Q2$ than in nucleotide. (b) The plot shows the normalized edit distance comparison for nucleotide vs $Q3$ in the 2D reads experiment setting. The slope of the regression line is 0.9208, indicating 7.92% well aligned reads have a better normalized edit distance in $Q3$ than in nucleotide.

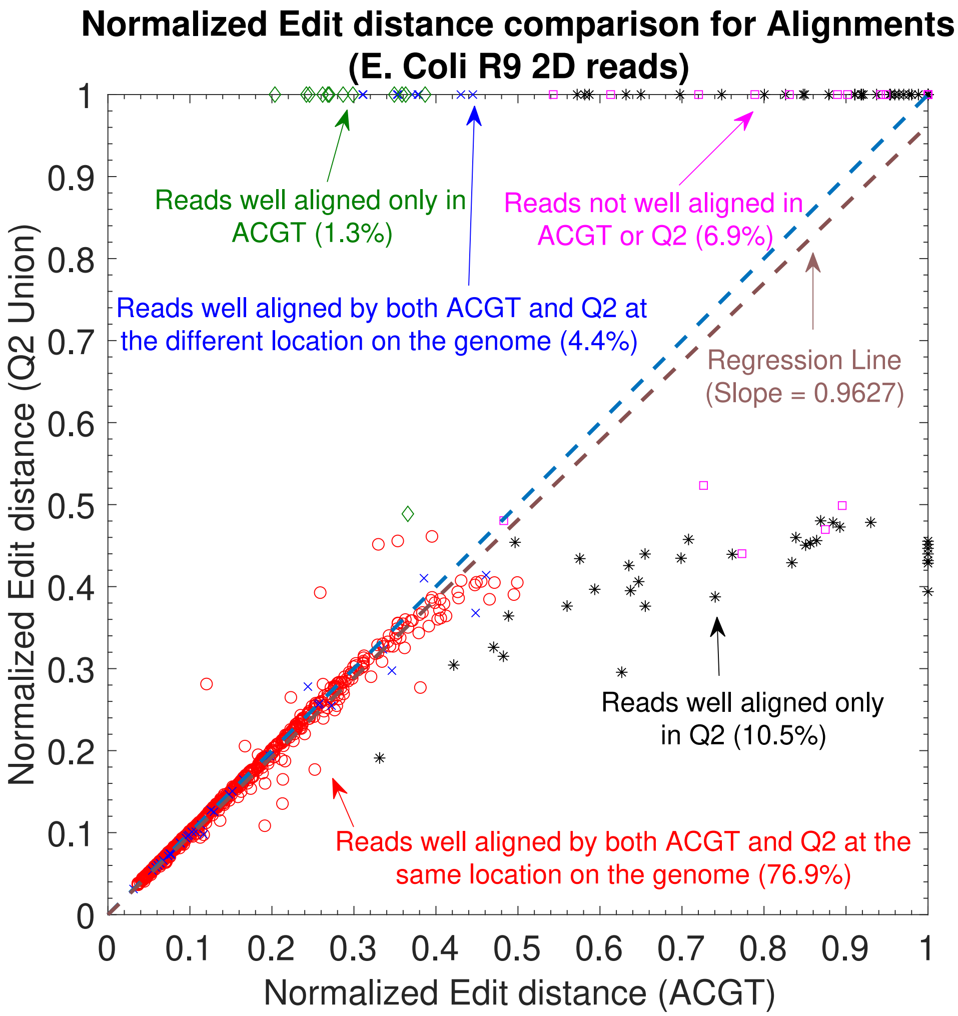


Figure 4: The alignment of long nanopore DNA-Seq R9 2D read onto E. Coli reference genome using the alternate approach that uses the information of the alignments of the corresponding 1D reads. The plot shows the normalized edit distance comparison for nucleotide vs $Q2$. The slope of the regression line is 0.9627, indicating 3.73% well aligned reads have a better normalized edit distance in $Q2$ than in nucleotide.

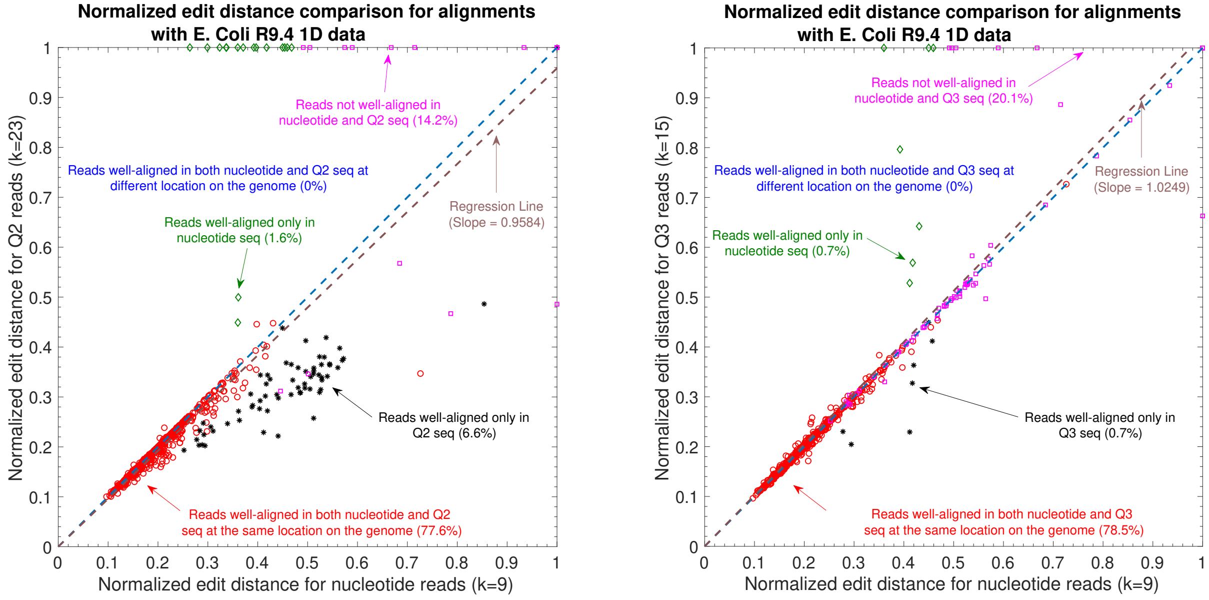


(a) (b)

Figure 5: The alignment of long nanopore DNA-Seq R9.4 1D read onto E. Coli reference genome (a) Normalized edit distance comparison for nucleotide vs $Q2$ (b) Normalized edit distance comparison for nucleotide vs $Q3$.

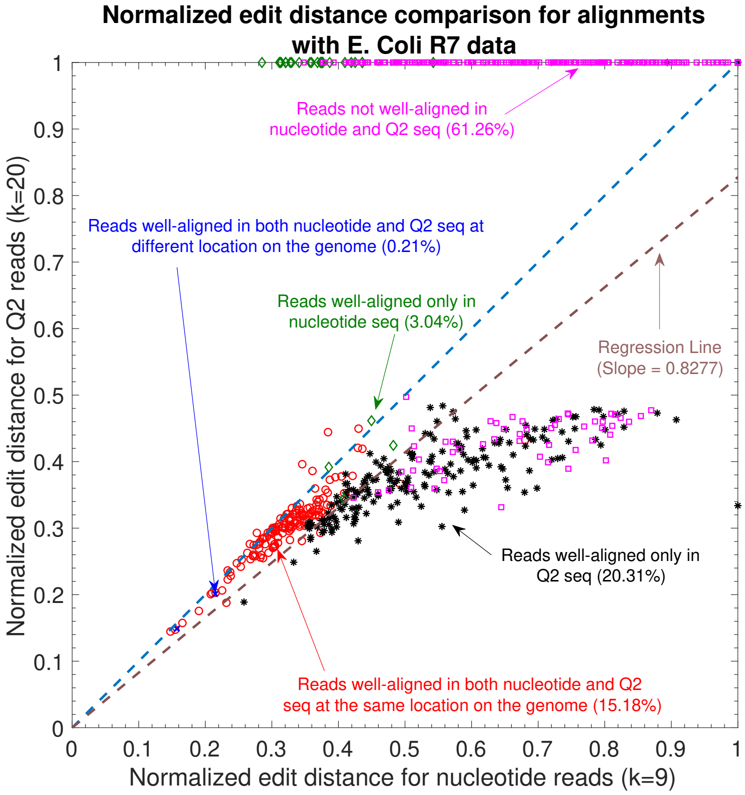


Figure 6: Normalized edit distance comaprison for nucleotide vs $Q2$ for E. Coli R7 1D reads. Majority of the reads are well-aligned by QAlign (35.7% of the 955 total reads are well-aligned in $Q2$, whereas only 18.43% reads are well-aligned in nucleotide). Moreover, the slope of the regression line is 0.8277 indicating an average gain of 17.23% in terms of the normalized edit distance.

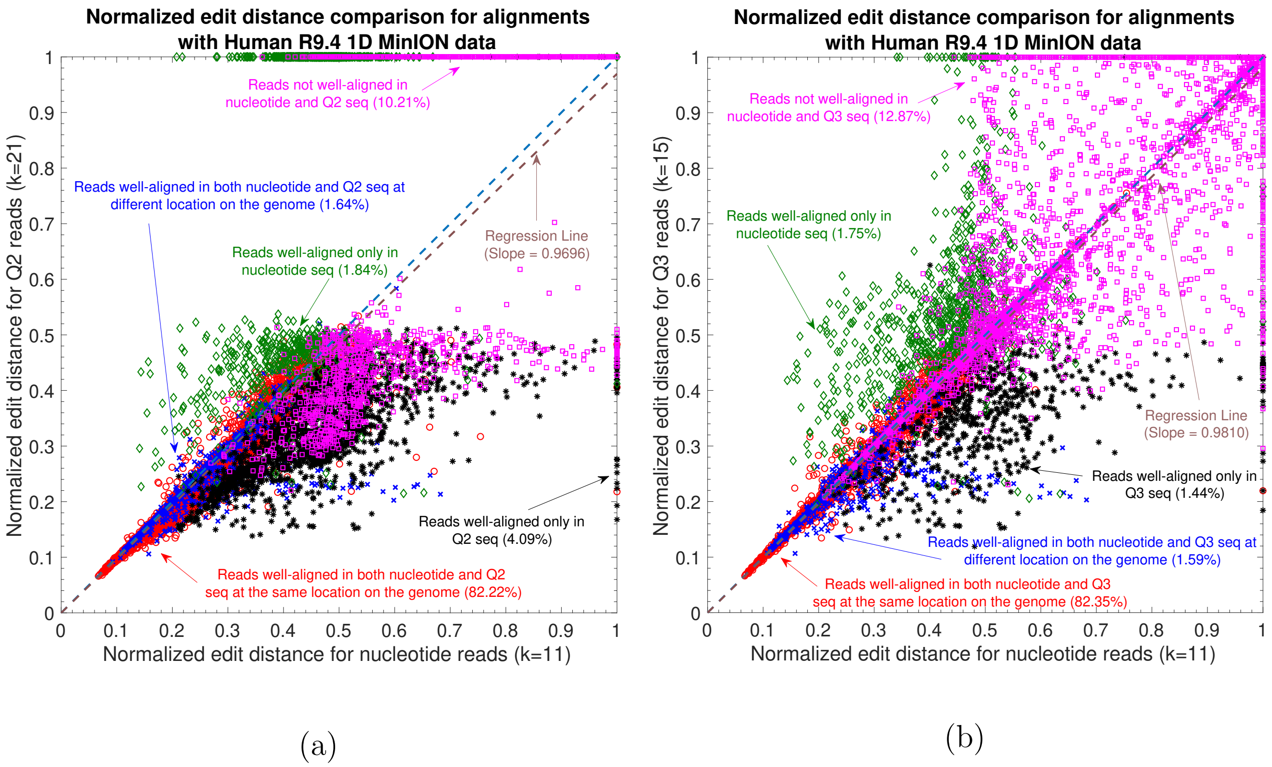


Figure 7: (a) Normalized edit distance comparison for read-to-genome alignments with $Q2$ and nucleotide Human R9.4 1D MinION data. The slope of the regression line is 0.9696, therefore, representing more accurate alignments in $Q2$. (b) Normalized edit distance for read-to-genome alignment with $Q3$ and nucleotide Human R9.4 1D MinION data. The slope of the regression line is 0.9810, therefore, representing more accurate alignments in $Q3$ as well.

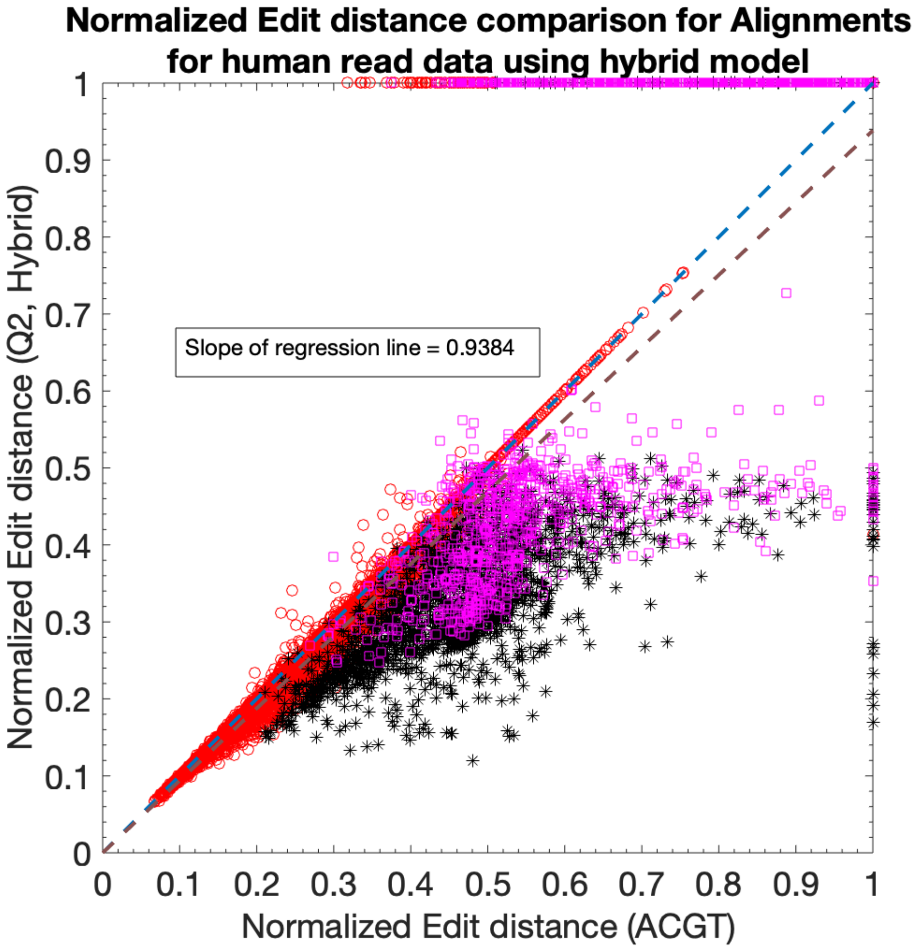


Figure 8: Normalized edit distance comparison for alignments using the hybrid model and the nucleotide alignments using the Human R9.4 1D MinION data. The hybrid model enhances the overall alignment accuracy by using nucleotide alignments when the $Q2$ alignments are missing and vice-versa. It is evident from the slope of the regression line as well as it decreases to 0.9384 from 0.9696 (without the hybrid model shown in Supplemental Figure 7a)

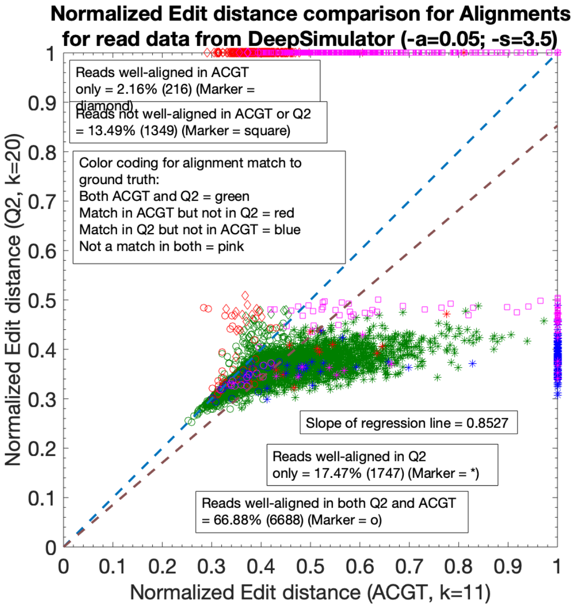


(a)

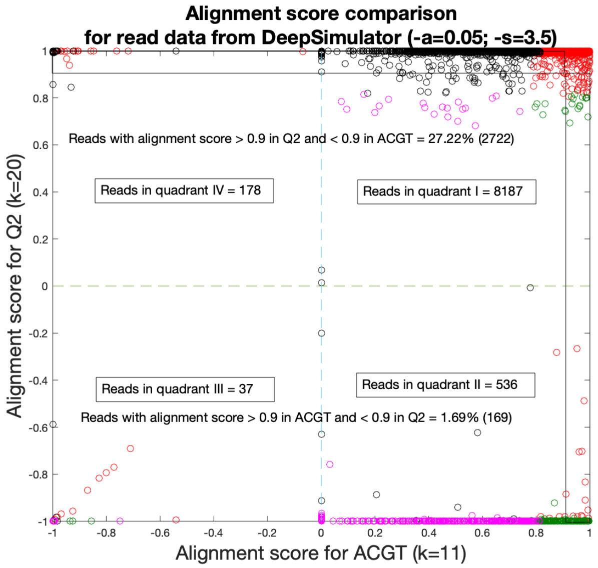


(b)

Figure 9: Benchmark for read-to-genome alignment using the simulated human reads from chromosome 1 of GRCh38 using Deep Simulator. (a) Normalized edit distance comparison for $Q2$ and nucleotide alignments using 10000 reads with an average read length of 12000. The Deep Simulator parameters used are corresponding to *low accuracy* reads, i.e., $a = 0.05$ and $s = 3.5$ using the context-dependent pore model. The color coding represents whether an alignment matches to the location in the ground truth - green represents both $Q2$ and nucleotide alignments matching in ground truth; red represents only nucleotide alignments matching in ground truth; blue represents only $Q2$ alignments matching in ground truth; pink represents neither of them matching the ground truth. The different markers represent if the reads are well-aligned in nucleotide or $Q2$ alignments as shown in the plot. (b) Alignment score comparison plot for $Q2$ and nucleotide alignments. For a known ground truth, we define alignment score as the ratio of the intersection of the alignment in the ground truth and the algorithm to the union of the alignment in the ground truth and the algorithm. It is computed similarly as *overlap quality* for read-to-read alignment. The negative score represents that the alignment by the algorithm has zero overlap with the alignment in the ground truth (mis-alignment).

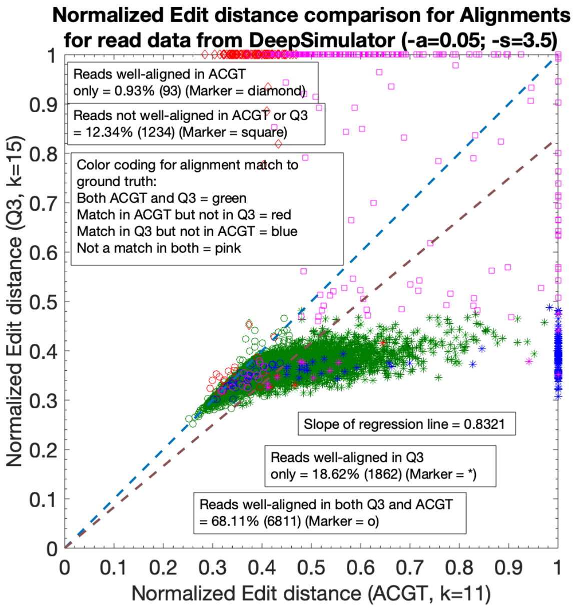


(a)

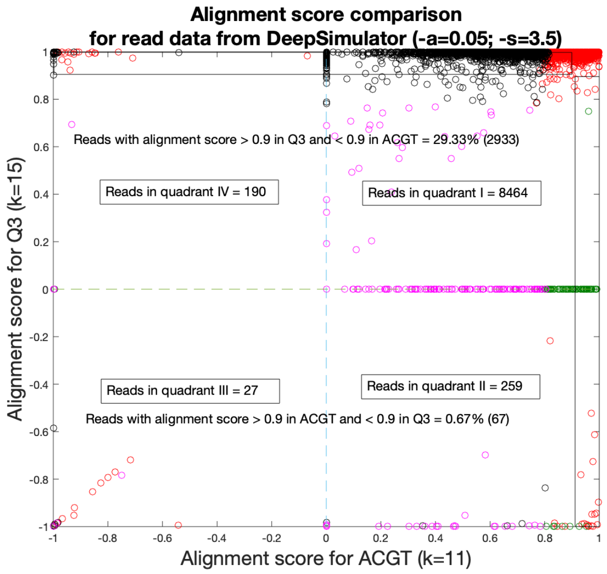


(b)

Figure 10: Benchmark for read-to-genome alignment using the simulated human reads from chromosome 1 of GRCh38 using Deep Simulator. (a) Normalized edit distance comparison for $Q2$ and nucleotide alignments using 10000 reads with an average read length of 12000. The Deep Simulator parameters used are corresponding to *low accuracy* reads, i.e., $a = 0.05$ and $s = 3.5$ using the context-dependent pore model. The color coding represents whether an alignment matches to the location in the ground truth - green represents both $Q3$ and nucleotide alignments matching in ground truth; red represents only nucleotide alignments matching in ground truth; blue represents only $Q3$ alignments matching in ground truth; pink represents neither of them matching the ground truth. The different markers represent if the reads are well-aligned in nucleotide or $Q3$ alignments as shown in the plot. (b) Alignment score comparison plot for $Q3$ and nucleotide alignments. For a known ground truth, we define alignment score as the ratio of the intersection of the alignment in the ground truth and the algorithm to the union of the alignment in the ground truth and the algorithm. It is computed similarly as *overlap quality* for read-to-read alignment. The negative score represents that the alignment by the algorithm has zero overlap with the alignment in the ground truth (mis-alignment).

#### 2 Read-to-Read alignment

*Methodology and Performance Metrics:* The ability of QAlign to align DNA reads to other DNA reads is discussed here. In each experiment, the DNA reads are aligned to other DNA reads using both QAlign and Minimap2. The read datasets from the same organisms are used for the experiments that we have used for read-to-genome alignment. We say that the algorithm is able to align the pair of reads (*i.e.*, the pair of reads has an overlap), if the 'Mapped Region' of overlap by the algorithm is atleast 90% of the estimated overlap between the same pair of reads in the ground truth (Please refer to the Method section 2.2.2 of the main paper for more details). Since we do not have a ground truth specifying what pairs of reads are aligning with each other with a head-to-tail alignment, therefore, in our evaluation we have used the read-to-genome alignments to leverage this information to estimate the ground truth for the read-to-read alignment. The overlap length between a pair of overlapping reads can be determined from the location of the alignments of the reads on the genome. We define the overlap quality as the ratio of the size of the intersection between the overlap region estimate (from the alignment algorithm) and the true overlap (from the ground truth), to the size of the union between the overlap region estimate and the true overlap. Thus, the overlap quality is equal to 1 if and only if the overlap region estimate is the same as overlap in the ground truth. The expected overlap quality is the area under the complementary cumulative distribution curve of overlap quality (as shown in Figure 4b). This expected value is also referred to as average overlap quality in this paper. Moreover, an ensemble model chooses the best overlap between a given pair of reads using the information of the longest overlap in nucleotide and in $Q2$.

The Minimap2 command used for the alignment with nucleotide sequences is:

```
minimap2 -cx ava-ont -k 10 reads.fasta reads.fasta
```

The Minimap2 command used for the alignments with the quantized sequences is:

```
minimap2 -cx ava-ont -k 20 --for-only reads_q2.fasta reads_q2.fasta

minimap2 -cx ava-ont -k 20 --for-only reads_q2.fasta rc_reads_q2.fasta
```

The two separate commands used for the alignment with the quantized sequence also aims to *explicitly* account for the *template* and *reverse complement* strands of the reads, and the best alignment between a pair of reads is determined by the maximum overlap length.

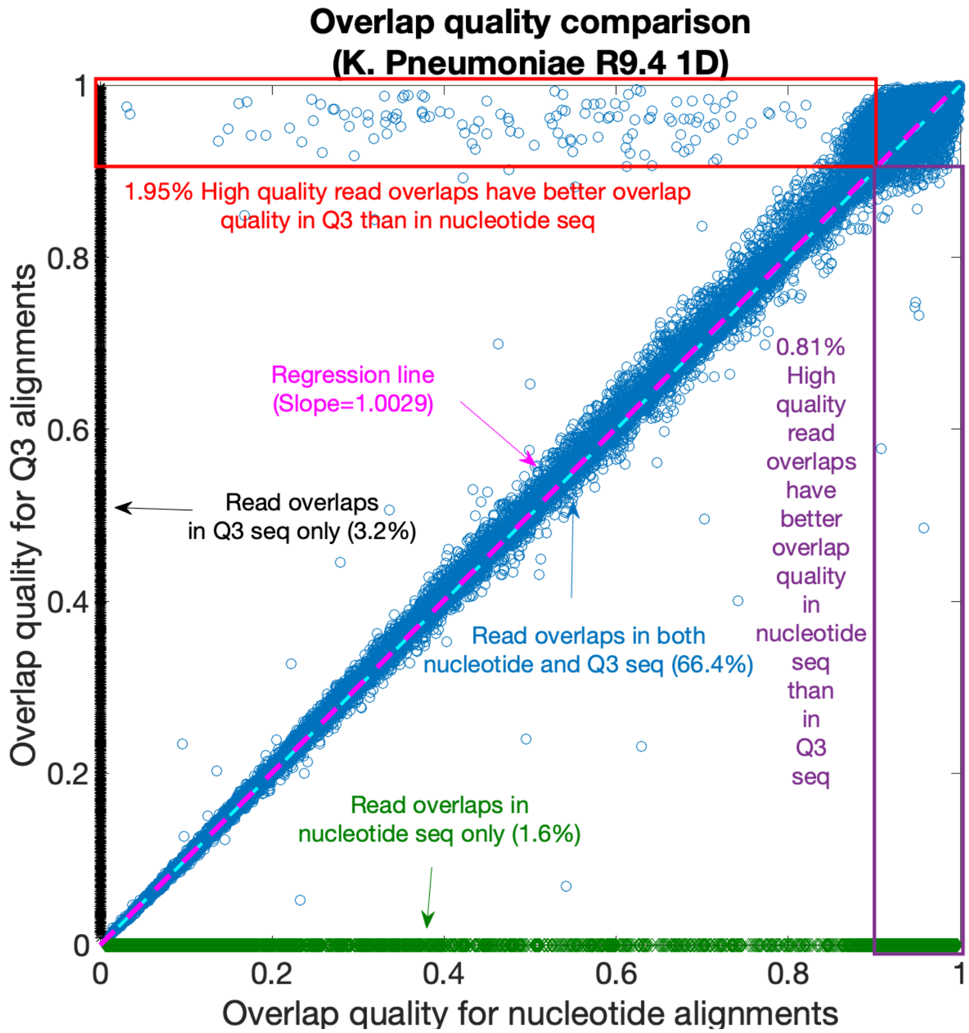


Figure 11: The read-to-read alignment of long nanorepore DNA-Seq reads (Overlap quality comaprison for nucleotide vs $Q3$ for K. Pneumoniae R9.4 1D dataset).

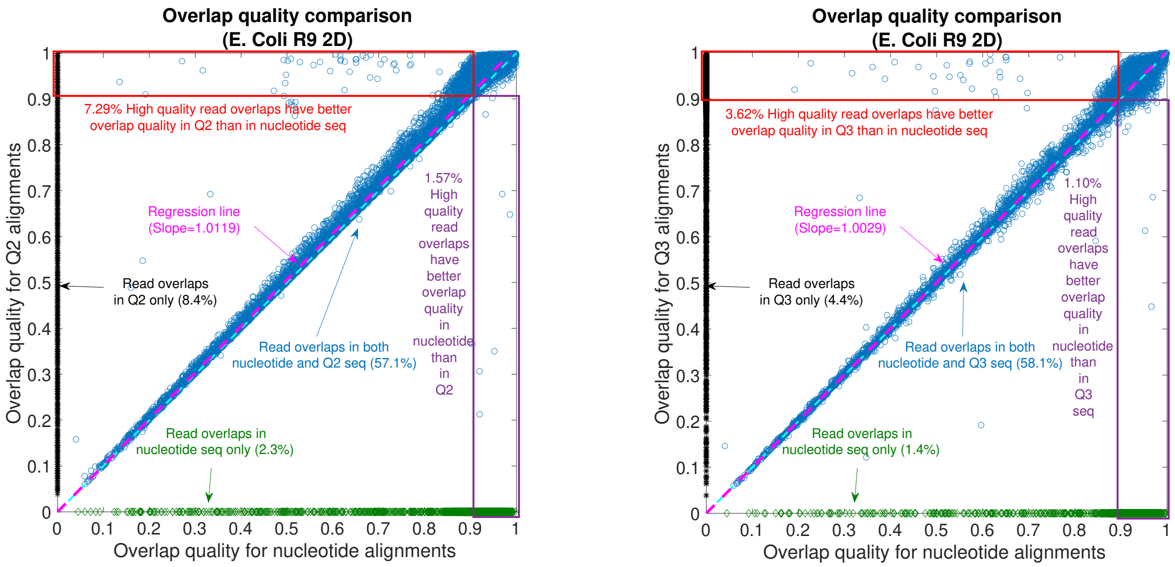


(a) (b)

Figure 12: The read-to-read alignment of long nanorepore DNA-Seq reads of E. Coli sequenced from MinION R9 2D flow cell. (a) Overlap quality comaprison for nucleotide vs $Q2$ alignments (b) Overlap quality comparison for nucleotide vs $Q3$ alignments.

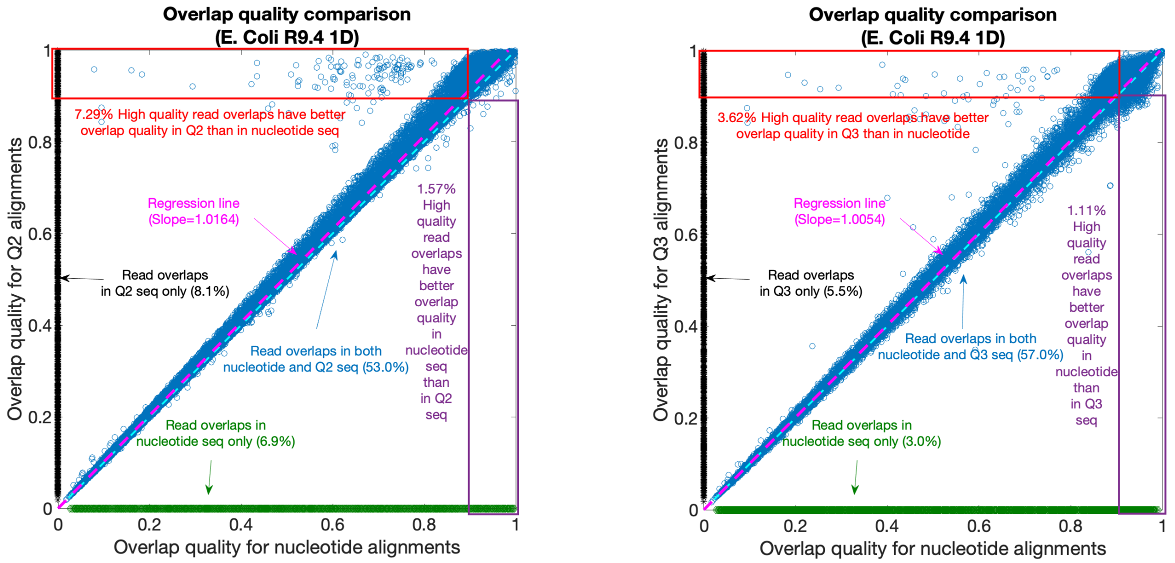


(a) (b)

Figure 13: The read-to-read alignment of long nanorepore DNA-Seq reads of E. Coli sequenced from MinION R9.4 1D flow cell. (a) Overlap quality comparison for nucleotide vs $Q2$ alignments (b) Overlap quality comparison for nucleotide vs $Q3$ alignments.

*Ensemble model for read-to-read alignment:* In case of the read-to-read alignment, there are several read overlaps that have better overlap quality in QAlign than in nucleotide alignment and several other read overlaps that have better overlap quality in nucleotide alignment than in QAlign. Therefore, in order to increase the *sensitivity*, an ensemble model is used, and the overlap quality comparison for the ensemble model against the nucleotide read overlaps is shown in Supplemental Figure 14.

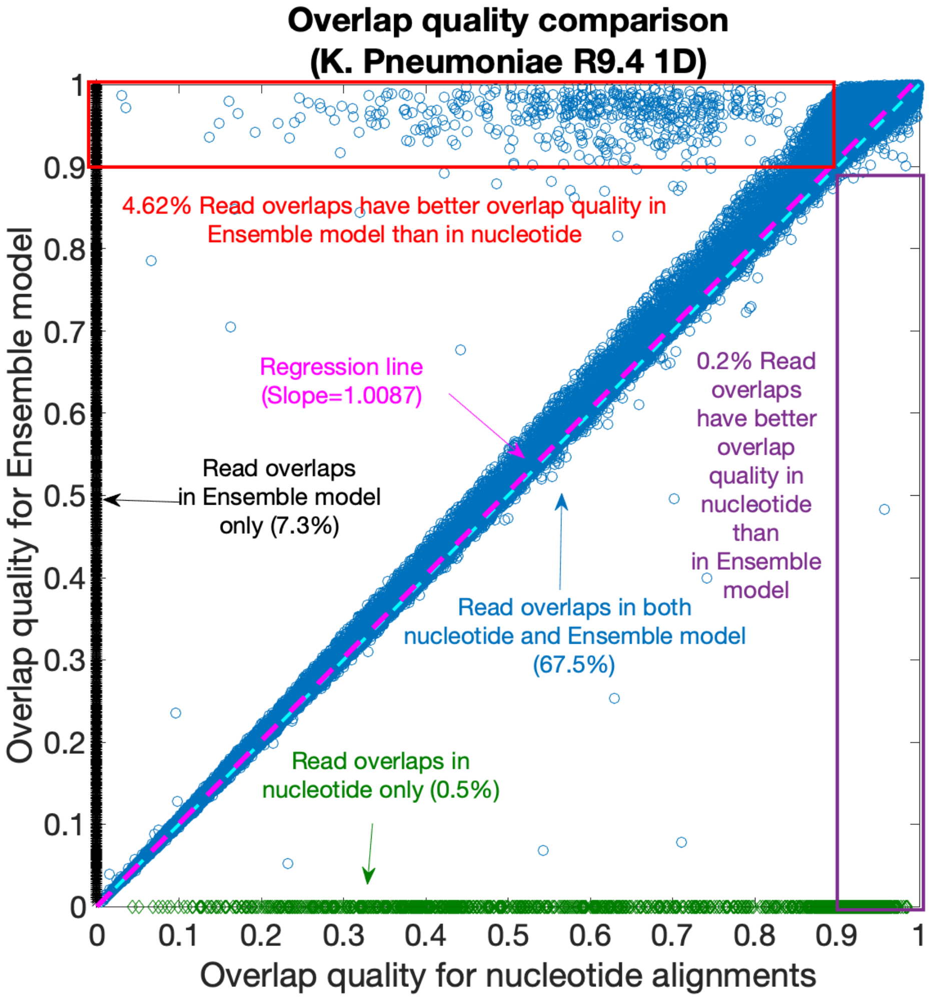


Figure 14: Ensemble model for read-to-read alignment for K. Pneumoniae R9.4 1D reads. We increase the *sensitivity* by capturing the alignments that are detected by either QAlign and nucleotide reads. In case of the common overlaps that are detected by both the algorithms, the one that has a longer overlap is chosen for the evaluation in this plot.

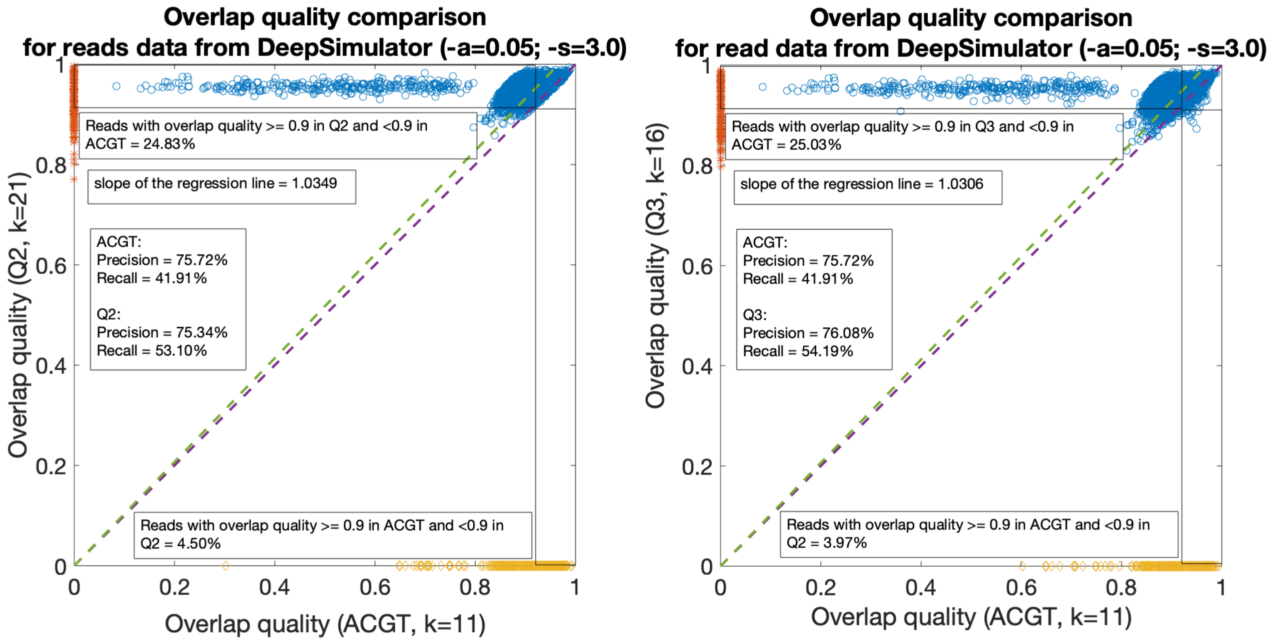


(a) (b)

Figure 15: (a) Overlap quality comparison for $Q2$ and nucleotide read-to-read alignment using simulated Human reads from Deep Simulator. (b) Overlap quality comparison for $Q3$ and nucleotide read-to-read alignment using simulated Human reads from Deep Simulator.

### 3 Read-to-Transcriptome alignment

*Methodology and Performance Metrics:* The methodology and the performance metrics are similar to the DNA read-to-genome alignment experiments: we compare the fraction of well-aligned reads as well as the normalized alignment length and the normalized edit distance for each alignment algorithm.

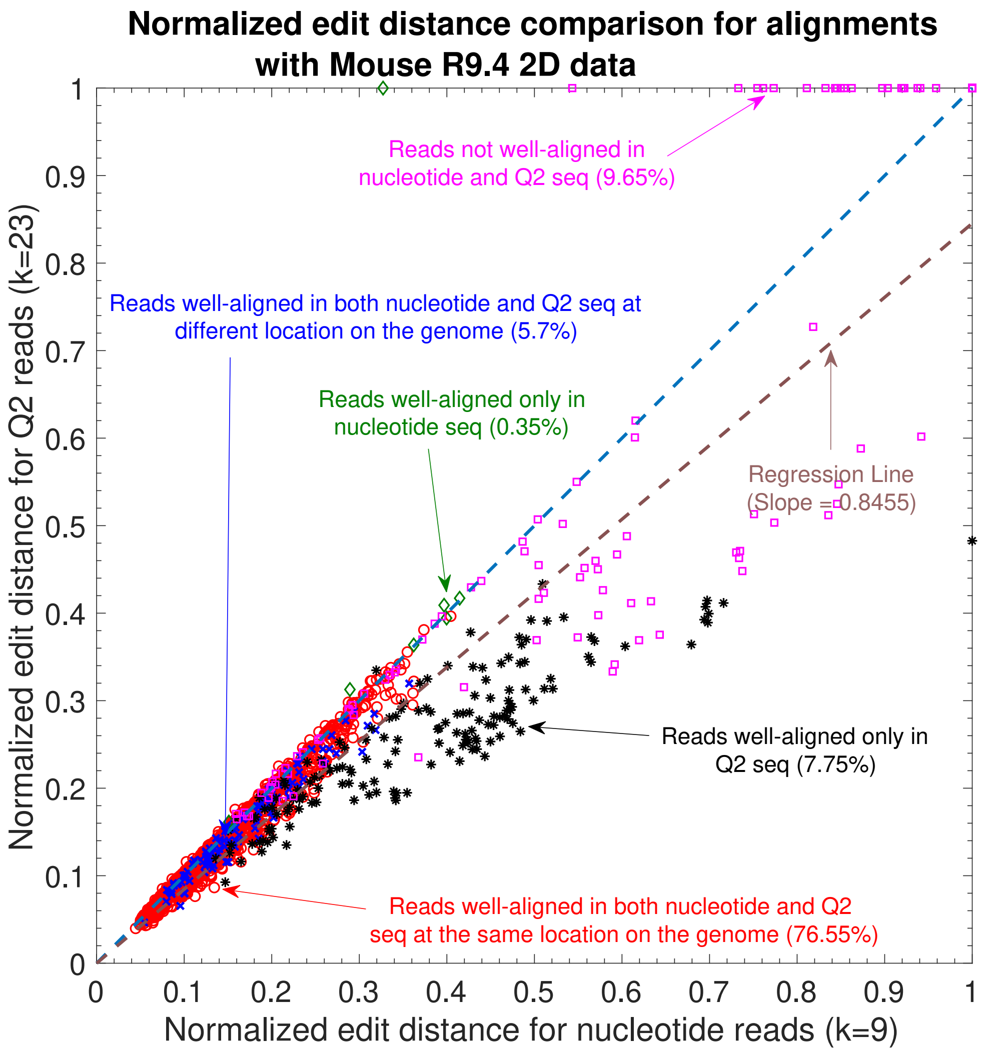


Figure 16: Normalized edit distance comparison for nucleotide vs $Q2$ alignments for Mouse R9.4 2D reads.

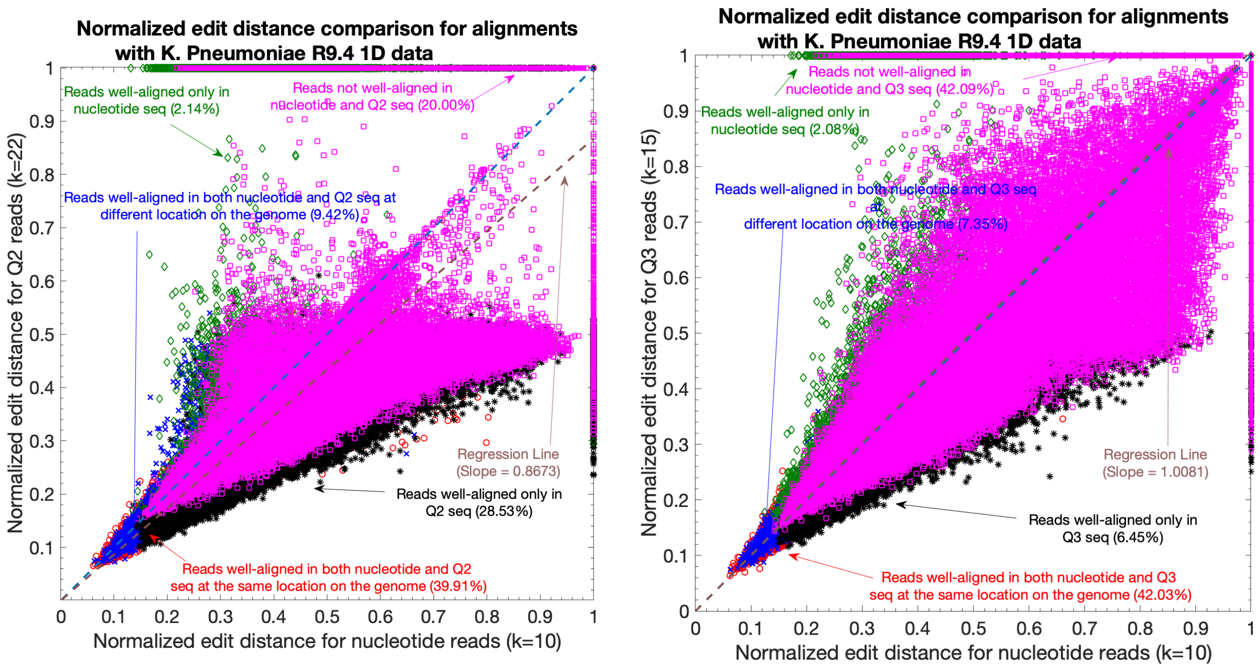


(a) (b)

Figure 17: (a) Normalized edit distance comparison for $Q2$ and nucleotide read-to-transcriptome alignment using 1 Million Human cDNA reads data. (b) Normalized edit distance comparison for $Q3$ and nucleotide read-to-transcriptome alignment using 1 Million Human cDNA reads data.

### 4 Normalized Edit Distance between Random DNA Sequences

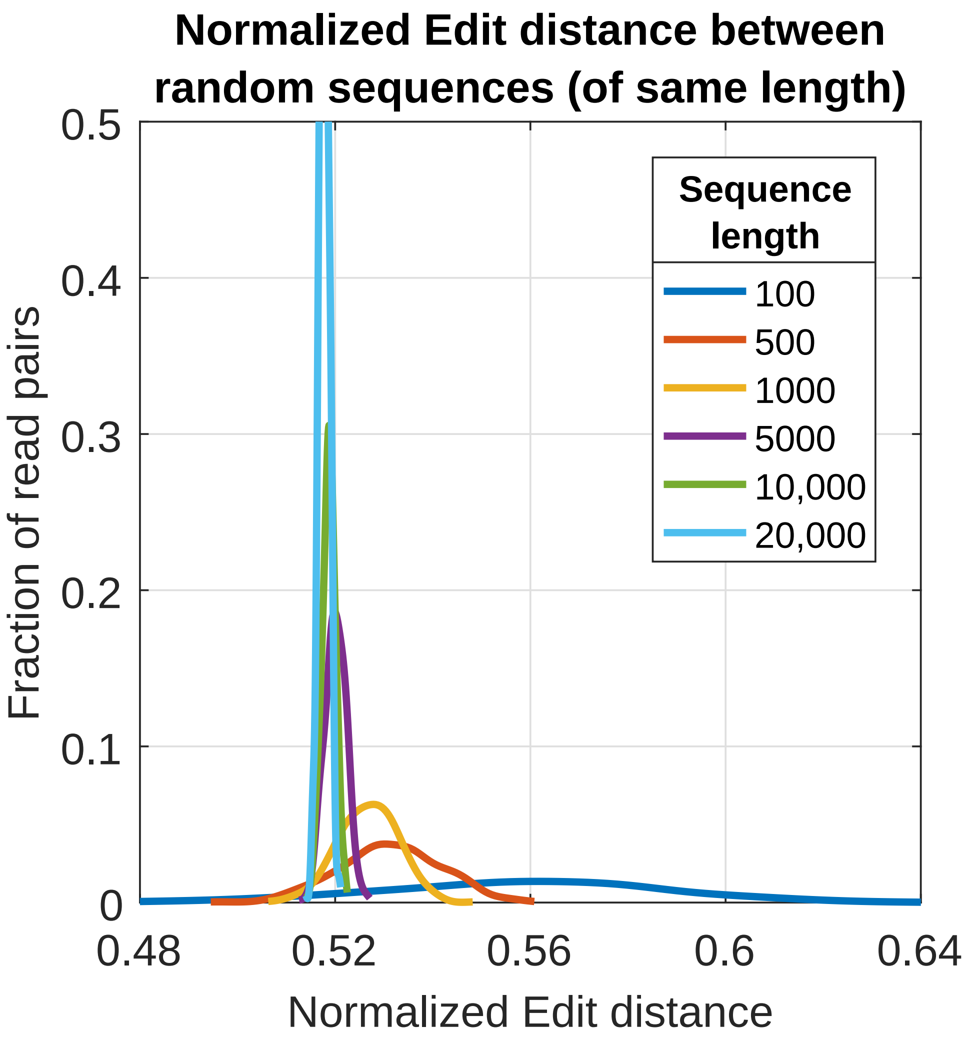


Figure 18: Empirical distribution for the normalized edit distance between pair of random DNA sequences. A random sequence has its symbols drawn i.i.d. from an alphabet e.g. $\Sigma = \{A, C, G, T\}$. We are interested in determining the minimum value of the normalized edit distance between two random sequences of same length, which can be used as a threshold to distinguish the alignment between a pair of random DNA sequences to pair of sequences that actually aligns to each other. We observe that the minimum value is 0.48, which we have used to comment if an alignment between a pair of DNA sequences is better (if the normalized edit distance is less than 0.48) than the alignment between a pair of random DNA sequences.

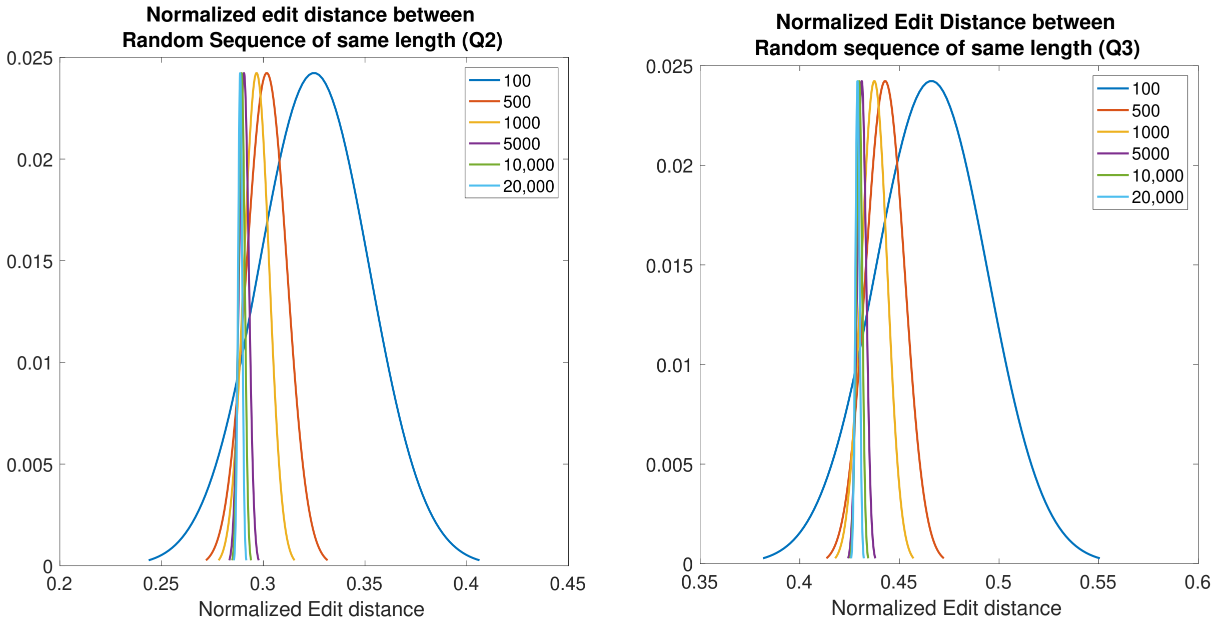


(a) (b)

Figure 19: (a) Empirical distribution for the normalized edit distance between pair of random binary sequences. A random binary sequence has its symbols drawn i.i.d. from an alphabet e.g. $\Sigma = \{0, 1\}$. We are interested in determining the minimum value of the normalized edit distance between two random binary sequences of same length, which can be used as a threshold to differentiate the alignment between a pair of random binary sequences. We observe that the minimum value is 0.25, which we have used to comment if an alignment between a pair of binary (or $Q2$) sequences is better (if the normalized edit distance computed by Minimap2 is less than 0.25) than the alignment between a pair of random binary sequences. (b) Empirical distribution for the normalized edit distance between pair of random ternary sequences. A random ternary sequence has its symbols drawn iid from an alphabet e.g. $\Sigma = \{0, 1, 2\}$. We observe that the minimum value is 0.35, which we have used to comment if an alignment between a pair of ternary (or $Q3$) sequences is better (if the normalized edit distance computed by Minimap2 is less than 0.35) than the alignment between a pair of random ternary sequences.

### 5 Methods

#### 5.1 Nanopore Sequencer

The Oxford Nanopore Technology is based on DNA transmigrated through a nanopore, which results in the changes in the ionic currents through the pore [1]. The current changes are caused by the different nucleotides that are partially blocking the pore as the (single-strand) DNA sequence to be measured is migrated through the nanopore. An enzyme slows down the motion of the DNA through the pore, so that the variations in the current signals can be measured accurately [1]. Base calling algorithms [2] are developed to infer the nucleotide sequence (A,C,G,T) from the measured current changes.

Ideally, it is desired that the DNA is migrated through the nanopore at a constant rate and the current signal recorded at a given time is only affected by a single nucleotide in the nanopore. Consequently, the DNA sequence can be decoded unambiguously with high probability.

However, in reality, there are several non-idealities due to the physics of the nanopore and the enzyme. (i) *Inter-symbol interference:* Since the nanopore is bigger than the size of a single nucleotide, the observed current at a given time is influenced by multiple (neighbor) bases or $Q$-mer (where $Q$ could be 4,5, or 6). (ii) *Random dwelling time:* The amount of time spent by each $Q$-mer of the DNA sequence in the nanopore may vary. So the rate at which the DNA sequence migrates through the nanopore is a stochastic process. (iii) *Segment insertion and deletion:* There are segments that are repeated as well as segments that migrate through the nanopore without registering a current reading. This results in redundant and missing segments. (iv) *$Q$-mer map fading:* The measured current level is a function of the corresponding $Q$-mer in the nanopore. However, this function is not deterministic, i.e., for the same $Q$-mer dwelling in the nanopore at a different time may produces a current level with some variations. This is also reflected in the $Q$-mer map (Figure 1c in main paper), where we have plotted the median value of the current levels along with the variances as the error-bar. (v) *Noisy samples:* Each observed current level is also subject to a random noise [1].

Table 1: This table shows the computation time trade-off with different choice of the minimizer length `k` for the read-to-genome alignment

| Dataset | Method of alignment | Minimizer length (`k`) | Computation Time (in seconds) | Percentage of well-aligned reads |
|---|---|---|---|---|
| K. Pneumoniae R9.4 1D | Nucleotide | 9 | 164.75 | 76.20 |
| | | 10 | 22.91 | 78.00 |
| | | 11 | 7.97 | 79.00 |
| | Q2 | 22 | 93.92 | 88.50 |
| | | 23 | 43.01 | 88.70 |
| | | 24 | 22.99 | 88.50 |
| | Q3 | 14 | 67.54 | 79.20 |
| | | 15 | 24.37 | 80.70 |
| | | 16 | 12.35 | 81.00 |
| E. Coli R9.4 1D | Nucleotide | 9 | 23.02 | 79.20 |
| | | 10 | 5.11 | 79.00 |
| | | 11 | 3.11 | 78.70 |
| | Q2 | 22 | 29.55 | 84.50 |
| | | 23 | 13.2 | 84.20 |
| | | 24 | 6.79 | 83.60 |
| | Q3 | 14 | 17.57 | 79.10 |
| | | 15 | 6.02 | 79.20 |
| | | 16 | 4.42 | 78.90 |
| E. Coli R9 2D | Nucleotide | 9 | 25.17 | 80.70 |
| | | 10 | 6.25 | 80.10 |
| | | 11 | 3.64 | 79.30 |
| | Q2 | 22 | 34.39 | 84.40 |
| | | 23 | 16.65 | 84.10 |
| | | 24 | 10.76 | 82.70 |
| | Q3 | 14 | 23.86 | 81.50 |
| | | 15 | 8.81 | 80.90 |
| | | 16 | 6.01 | 80.10 |

Table 2: This table shows the computation time trade-off with different choice of the minimizer length k for the read-to-read alignment

| **Dataset** | **Method of alignment** | **Minimizer length (k)** | **Computation Time (in seconds)** | **Percentage of well-aligned reads** |
|---|---|---|---|---|
| K. Pneumoniae R9.4 1D | Nucleotide | 10 | 4118.63 | 68.07 |
| | | 11 | 2388.37 | 67.00 |
| | | 12 | 2263.17 | 66.00 |
| | Q2 | 21 | 4864.79 | 71.98 |
| | | 22 | 4477.58 | 70.92 |
| | | 23 | 4425.29 | 69.73 |
| | Q3 | 14 | 3715.9 | 69.60 |
| | | 15 | 2854.07 | 68.73 |
| | | 16 | 2768.73 | 67.77 |
| E. Coli R9.4 1D | Nucleotide | 11 | 882.78 | 62.27 |
| | | 12 | 824.00 | 60.49 |
| | | 13 | 808.74 | 58.72 |
| | Q2 | 21 | 1169.15 | 62.60 |
| | | 22 | 1134.86 | 60.82 |
| | | 23 | 1115.41 | 58.87 |
| | Q3 | 14 | 985.57 | 63.87 |
| | | 15 | 914.89 | 62.53 |
| | | 16 | 903.19 | 60.92 |
| E. Coli R9 2D | Nucleotide | 10 | 727.56 | 59.99 |
| | | 11 | 560.08 | 58.78 |
| | | 12 | 510.38 | 57.78 |
| | Q2 | 20 | 1129.05 | 65.47 |
| | | 21 | 1074.90 | 64.82 |
| | | 22 | 1048.20 | 64.14 |
| | Q3 | 14 | 705.86 | 62.46 |
| | | 15 | 640.17 | 62.00 |
| | | 16 | 621.43 | 61.44 |

Table 3: This table shows the computation time trade-off with different choice of the minimizer length `k` for the read-to-genome alignment of 50000 Human R9.4 1D reads.

| **Method of alignment** | **Minimizer length (`k`)** | **Computation Time (in seconds)** | **Percentage of well-aligned reads** |
|---|---|---|---|
| Nucleotide | 11 | 326,270 | 85.69 |
| | 12 | 38,446 | 85.11 |
| | 13 | 12,030 | 84.36 |
| | 14 | 4100 | 83.69 |
| | 15 | 2202 | 82.89 |
| Q2 | 21 | 268,864 | 87.94 |
| | 22 | 149,614 | 87.45 |
| | 23 | 68,403 | 86.87 |
| | 24 | 50,913 | 86.19 |
| | 25 | 44,028 | 85.49 |
| Q3 | 15 | 203,926 | 85.37 |
| | 16 | 63,994 | 85.10 |
| | 17 | 29,428 | 84.54 |
| | 18 | 16,182 | 84.07 |
| | 19 | 10,615 | 83.45 |

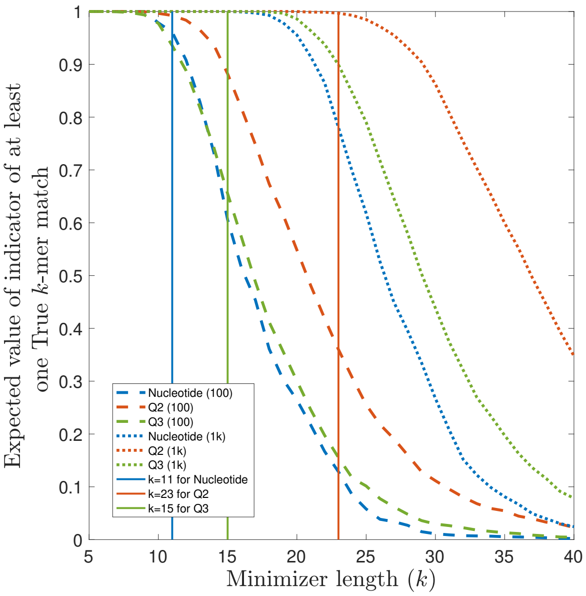


(a)

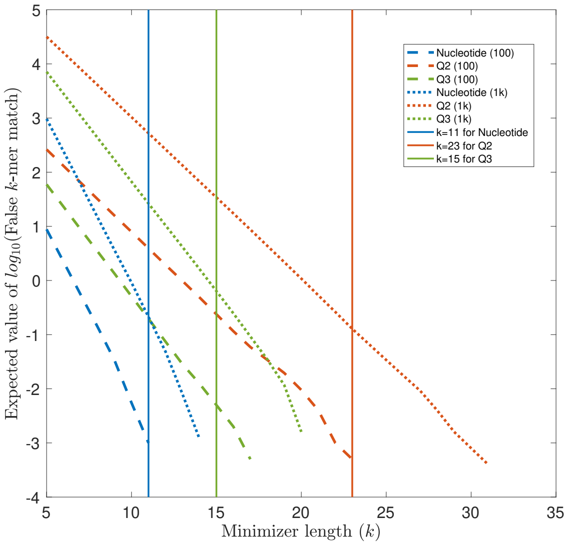


(b)

Figure 20: **Why short alignments are difficult with QAlign?** In this experiment, a random DNA sequence $s_1$ of length 100 and 1000 are passed through an i.i.d. substitution channel with an error rate or 15% and the output sequence from the channel is $s_2$. The corresponding quantized sequences are $s_1^Q$ and $s_2^Q$, respectively. A "True $k$-mer match" is a sub-sequence of length $k$ that matches exactly at the same position on $s_1$ and $s_2$ (or $s_1^Q$ and $s_2^Q$). A "False $k$-mer match is when there is an exact match of length $k$ but at different location on $s_1$ and $s_2$ (or $s_1^Q$ and $s_2^Q$). The indicator function returns 1 in case there is at least one "True $k$-mer match". The expected value of the indicator function gives an estimate of the probability of finding at least one "True $k$-mer match". It is evident from figure (a) that the Expected value for $Q2$ for a choice of minimizer length $k = 23$ is smaller than the expected value for Nucleotide for $k = 11$ when small sequence of length 100 is considered. Whereas the expected value for $Q2$ is nearly 1 for same choice of $k = 23$ when sequence of length 1000 is considered. On the other hand, choosing a smaller $k$ for $Q2$ (for example, $k = 11$ for $Q2$ which gives the expected value for True $k$-mer match to be nearly 1) leads to more "False $k$-mer matches", and hence, it requires more computation time as compared to Nucleotide alignments. Therefore, the alignments of small chunks of reads is difficult in QAlign, which is likely the case in spliced-alignments of RNA reads to genome or read-to-read alignment when the overlaps is as small as a few hundreds of bases.

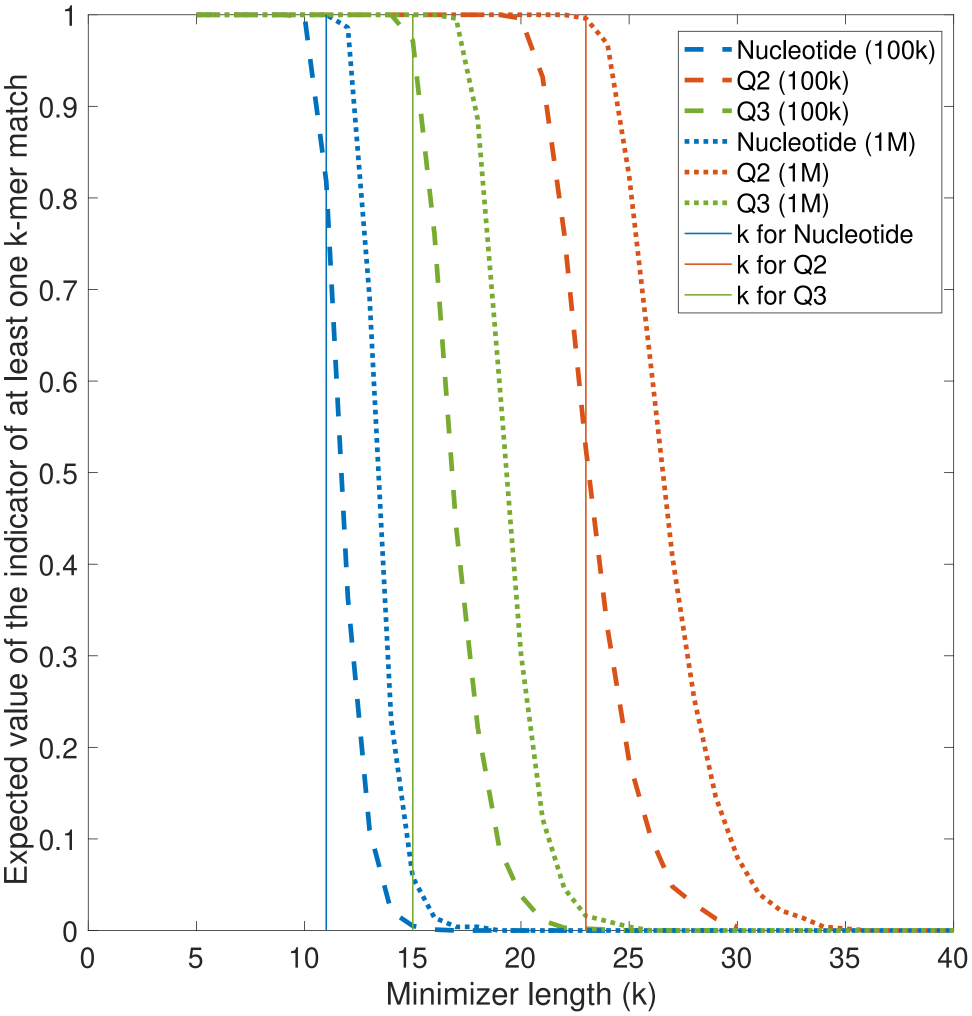


Figure 21: The plot shows the expected value of the indicator of at least one $k$-mer match when all the $k$-mers from a random DNA Genome (of length 100000 and 1000000) are matched against all the $k$-mers from a random DNA read (of length 1000). The expected value also represent the probability of at least one false $k$-mer match since both the genome and the read sequence are randomly generated. It shows that as the length of the genome increases the probability of a random $k$-mer match also increases for the same value of $k$.

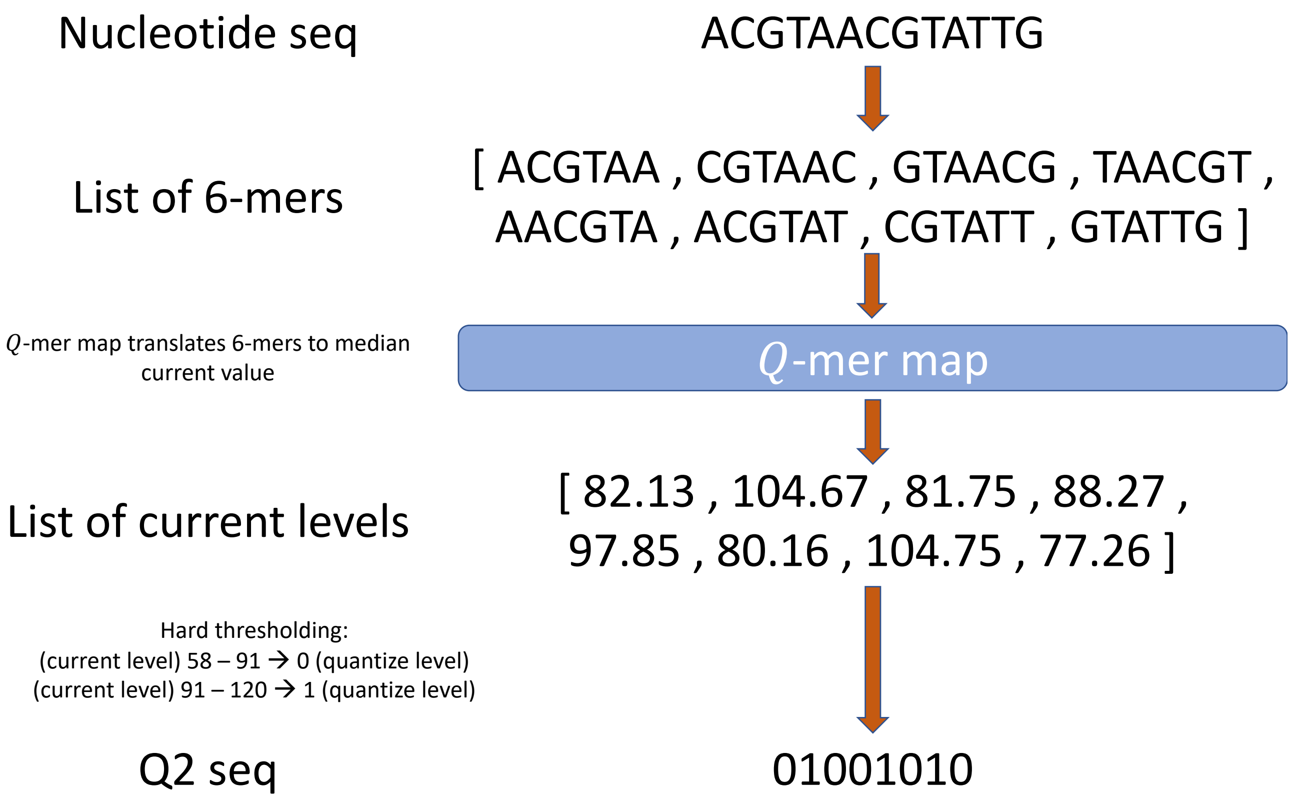


Figure 22: An example for the Quantization method for QAlign.

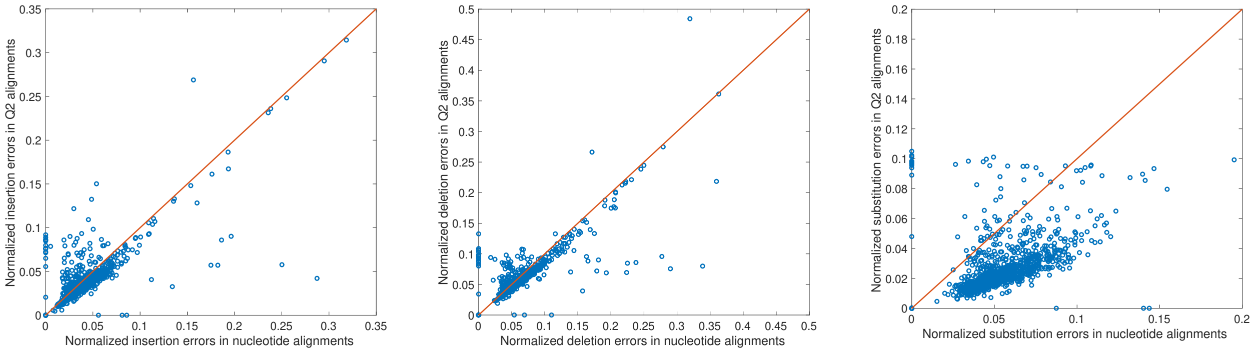


(a) (b) (c)

Figure 23: A comparison for the insertion, deletion, and substitution errors for nucleotide and $Q2$ alignments for K. Pneumoniae R9.4 1D reads dataset. The normalized errors are defined as number of errors normalized by the length of the alignment. For example, normalized insertion errors is computed as the ratio of total number of insertions in the alignment of a read to the length of the alignment. The comparison is provided for the 1000 reads used for the read-to-genome alignment. We observe that the normalized insertion and normalized deletion errors in $Q2$ are nearly same as that of nucleotide alignments. However, the normalized substitution errors in $Q2$ alignments is much low than that of nucleotide alignments across the 1000 reads.

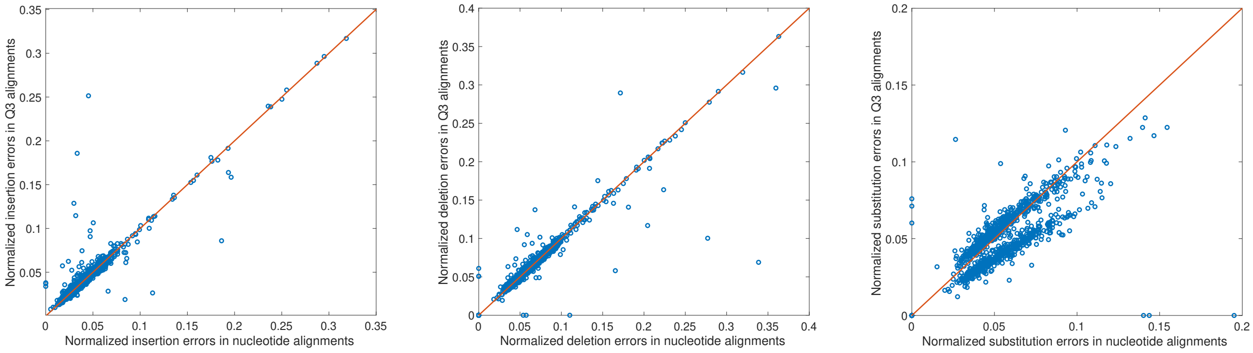


(a) (b) (c)

Figure 24: A comparison for the insertion, deletion, and substitution errors for nucleotide and $Q3$ alignments for K. Pneumoniae R9.4 1D reads dataset. The normalized errors are defined as number of errors normalized by the length of the alignment. For example, normalized insertion errors is computed as the ratio of total number of insertions in the alignment of a read to the length of the alignment. The comparison is provided for the 1000 reads used for the read-to-genome alignment. We observe that the normalized insertion and normalized deletion errors in $Q3$ are nearly same as that of nucleotide alignments. However, the normalized substitution errors in $Q3$ alignments is slightly lower than that of nucleotide alignments across the 1000 reads.
